# Unilateral ephaptic program underlying sweetness dominance

**DOI:** 10.1101/2023.08.04.551918

**Authors:** MinHyuk Lee, Seon Yeong Kim, Taeim Park, Kyeung Min Joo, Jae Young Kwon, Kyuhyung Kim, KyeongJin Kang

## Abstract

In ephaptic coupling, physically adjacent neurons influence one another’s activity via the electric fields they generate. To date, the molecular mechanisms that mediate and modulate ephaptic coupling’s effects remain poorly understood. Here, we show that the hyperpolarization-activated cyclic nucleotide-gated (HCN) channel lateralizes the potentially mutual ephaptic inhibition between *Drosophila* gustatory receptor neurons (GRNs). While sweet-sensing GRNs (sGRNs) engage in ephaptic suppression of the adjacent bitter-sensing GRNs (bGRNs), HCN expression in sGRNs enables them to resist ephaptic suppression from the bGRNs. Such one-sided ephaptic inhibition confers sweetness dominance, facilitating ingestion of bitter-laced sweets. Flies with HCN-deficient sGRNs exhibited dramatically decreased attraction to sucrose mixed with moderate levels of caffeine, highlighting the behavioral significance that gustatory ephaptic inhibition promotes ingestion of carbohydrates buried in bitterness. Our findings indicate a role for the gating of ephaptic coding to ensure the intake of the essential nutrient despite bitter contaminants present in the feeding niche of *Drosophila*, as the gating establishes a hierarchy of gustatory neuron excitation. Such refinement provides a previously unappreciated mechanism for controlling the activity of a neuronal network with potential implications in the mammalian brain, given the evolutionary conservation of the HCN genes.

## Introduction

Electrical activity in the form of action potentials mediates intraneuronal signal propagation. This can lead to extracellular potential changes that increase or decrease the excitability of adjacent neurons independent of synapses, a phenomenon called ephaptic coupling^1–3^. Although the mechanisms underlying ephaptic coupling and their implications are unclear, such coupling is widespread in the brain^4–7^ and thought to underlie the temporal synchronization of neuronal firing that contributes to generation of neural oscillations^4,5,8^. Such coupling can be dramatic when two neurons form complex interfaces like the pinceau formed between basket cells and Purkinje neurons in mice ^9^, the axon cap formed between PHP interneurons and Mauthner cells in fish ^10^, and those formed among the compartmentalized olfactory receptor neurons (ORNs) in *Drosophila* ^11–13^. All these cases are examples of ephaptic inhibition^3^. The latter two examples are characterized by bilateral inhibition between the participating neurons^12,14,15^. Interestingly, differences in cell size lead to asymmetric lateral suppression, with larger *Drosophila* ORNs dominating their smaller partners ^12^. Ephaptic inhibition occurs when excitation of a neuron hyperpolarizes a partner target neuron ^3,9,12^. We still understand little about the genetic programming and regulation of ephaptic coupling beyond its most basic features, such as structural complexity and hyperpolarization of the inhibited post-ephaptic neurons.

Carbohydrates, the primary energy source for animals, are produced by plants and other photosynthetic organisms. Sugar-producing organisms often deter predators with bitter compounds that warn animals of potential post-ingestive toxicity ^16^. To avoid the toxicity, sugar ingestion in *Drosophila* has been reported to be suppressed in the presence of bitters ^17–21^. This can be achieved by two distinct mechanisms, the odorant-binding protein OBP49a ^21^ and a GABA-mediated inhibitory circuit ^17^. At the same time, animals must overcome the bitter-induced aversion to feed on bitter-laced sugars as found in plants. Human gustation is largely tuned to favor sugar ingestion, with sweetness reducing the perception of other primary taste modalities in psychophysiological studies ^22–29^. Yet, little is known about the sensory processing mechanisms underlying sweetness dominance. In *Drosophila*, their phytophagy suggests the existence of a system for sugar-induced bitterness suppression to facilitate adequate carbohydrate intake, but, as in other animals, the mechanisms of sweetness dominance in *Drosophila* remain unclear. Here, we describe how ephaptic inhibition between sweet- and bitter-sensing gustatory receptor neurons (s- and bGRNs) is lateralized to enable sweet-dependent bitter taste suppression.

## Results

### Sucrose inhibits bitter sensing and avoidance

Serendipitously, we observed that extracellular recordings of labellar bristle sensilla with bitter/sucrose mixtures produced significantly smaller responses from bGRNs than bitter-only controls (Fig. 1A,B, and Fig. S1A,B). Both i- and s-type labellar bristle sensilla house a bGRN and an sGRN (Fig. 1A) ^30–32^. The spiking responses from the sensilla of WT (*w*^1118^ on a Canton-S background) in response to bitter/sweet mixtures were readily discernible into two categories based on spike amplitude in both the depolarization and repolarization phases (Fig. S1A). To control for any effect of osmolarity, the bitter-only control solution contained equimolar sorbitol, which is nonsweet to flies ^33^. Sucrose suppressed responses to various bitters that stimulate i- and s-type labellar bristle sensilla, indicating that sucrose-dependent inhibition is common to bGRNs regardless of bristle type (Fig. 1A-D, Fig. S1B). We found that glycerol, which stimulates sGRNs ^34^, also suppressed bGRN activity (Fig. S1C,D). Next, we examined whether there are behavioral effects analogous to the observation with the recordings. To this end, we used capillary feeder (CAFE) assays ^35,36^ in which flies were offered two sets of capillaries containing the same concentration of sucrose plus or minus a bitter compound. Thus, it was investigated whether bitter avoidance can be attenuated by increasing sucrose concentrations. We found increasing concentrations of sucrose reduced bitter avoidance (Fig. 1E-G), indicating that sweetness suppresses the feeding avoidance induced by bitter chemicals. This inhibitory effect was not due to increased osmolarity, however, because equi-osmolar substitutions of sucrose with nonsweet sorbitol restored the avoidance (Fig. 1F,G). Note that we previously found that 30 mM sucrose is the minimal condition to induce ingestion for examination of bitter avoidance in CAFE. Lower sucrose concentrations did not elicit sufficient capillary feeding, with which we were unable to reliably appraise feeding aversion ^36,37^. Therefore, tastant concentrations used for CAFE were different from the recording experiments and the effect of sucrose concentrations was compared to the baseline acquired at 30 mM sucrose. Nevertheless, the increasing sucrose concentrations reduce aversion to various bitters, similar to the electrophysiological findings.

**Fig. 1.**
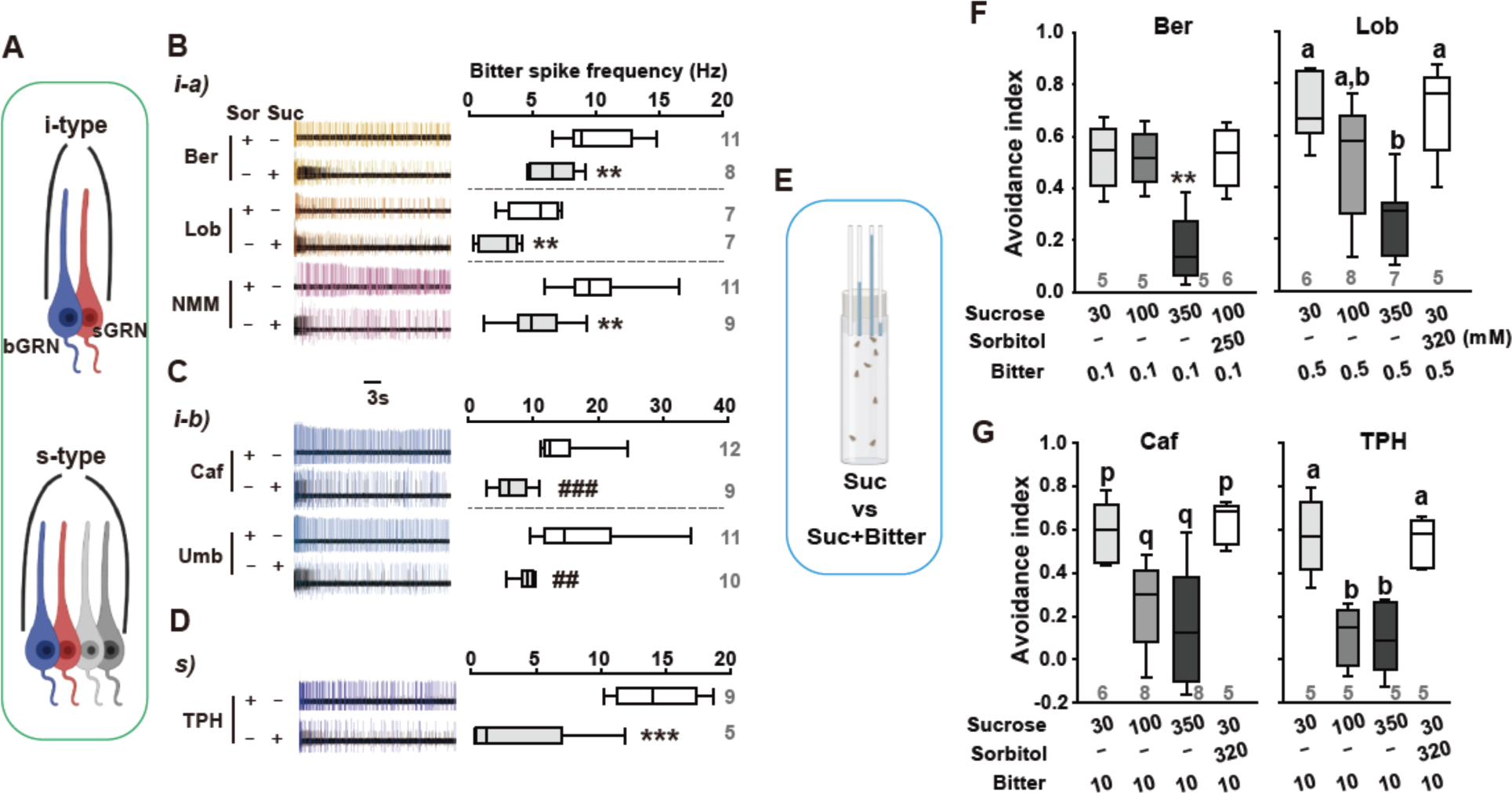
Sucrose suppresses bitter-sensing GRNs and ingestion avoidance of bitters. (**A**) Schematic showing GRN organization in sensilla of the indicated bristle type in the *Drosophila* labellum. Each sensillum harbors both a bitter- and sweet-sensing GRN (bGRN and sGRN). The s-type sensilla contain water- and salt-sensing GRNs (gray) in addition to s- and bGRNs. (**B-D**) Sucrose-induced bitter suppression in i-a (B), i-b (C), and s-type (D) bristle sensilla. Spikes from sweet-sensing GRNs (sGRNs) are black, while those from bGRNs are colored (left). See also Figure S1. Box plots showing the spike frequency distributions for the first 5 sec. Sor: sorbitol, 50 mM; Suc: sucrose, 50 mM (Sor and Suc were 10 mM for TPH); Ber: berberine, 0.5 mM; Lob: lobeline 0.5 mM; NMM: N-methyl maleimide, 2 mM; Caf: caffeine, 2 mM; Umb: umbelliferone, 0.5 mM; TPH: theophylline, 5 mM. (**E**) Schematic showing the capillary feeder assay. (**F** and **G**) Ingestion avoidance of bitters is suppressed by sucrose. **, ***: Student’s or Tukey’s t-test, p < 0.01, 0.001, respectively. ##, ###: Mann-Whitney U test, p < 0.01, 0.001, respectively. Letters indicate statistically distinct groups (p and q: Dunn’s test, and a and b: Tukey’s test, p < 0.05). Numbers in gray indicate the number of experiments.

### Sucrose-induced bitterness suppression requires sGRN excitation but not chemical synaptic transmission

Next, we examined the mechanism of sucrose-induced inhibition of bGRNs, testing whether the inhibition of bGRNs requires sucrose sensing by the sGRNs. The *Gr64af* deletion mutant lacks the entire *Gr64* gene locus from *Gr64a* to *Gr64f* and cannot detect sucrose^34^. In the i- and s-type sensilla of *Gr64af* mutants, the bitter/sucrose mixture elicited spiking responses only from bGRNs. We found that the addition of sucrose to a bitter tastant elicited a similar bGRN response as the bitter-only control (Fig. 2A, Fig. S2A). In CAFE assays, increasing concentrations of sucrose did not produce any significant suppression of bitter avoidance in *Gr64af* mutants (Fig. 2B). We next used *Gr5a-Gal4* ^38^ to drive the overexpression of the inwardly rectifying potassium channel Kir2.1^39^ to silence sGRNs with temporal control provided by *Gal80^ts^* (Fig. 2C) ^40^. When we activated the silencing in adults by raising the temperature to 29°C, the sucrose-induced inhibition of caffeine-activated bGRNs was abolished (Fig. 2C, Fig. S2B). In contrast, blocking chemical presynaptic transmission in sGRNs by similarly expressing the tetanus toxin light chain (TNT)^41^ did not relieve sucrose-dependent bGRN inhibition (Fig. 2D, Fig. S2C). We then confirmed these electrophysiological findings in CAFE assays, where blocking action potentials, but not chemical synaptic transmission, allowed the persistence of caffeine avoidance even in the presence of 350 mM sucrose (Fig. 2E). We next used RNAi in bGRNs to knock down the inhibitory neurotransmitter receptor genes *rdl* ^42^ and *gluClα* ^43^, which mediate fast GABAergic and glutamatergic postsynaptic inhibition, respectively. Additionally, in sGRNs, we targeted *shakB,* which encodes a major neuronal gap junction channel ^44^, although neurons coupled with electrical synapses share the direction of activation unlike the GRN inhibition we observed here. None of these knock-down experiments resulted in significant electrophysiological impairment of the lateral inhibition between GRNs (Fig. S3). These results suggest that peripheral sGRN excitation, and not synaptic feedback to bGRNs, is required for the bGRN inhibition.

**Fig. 2.**
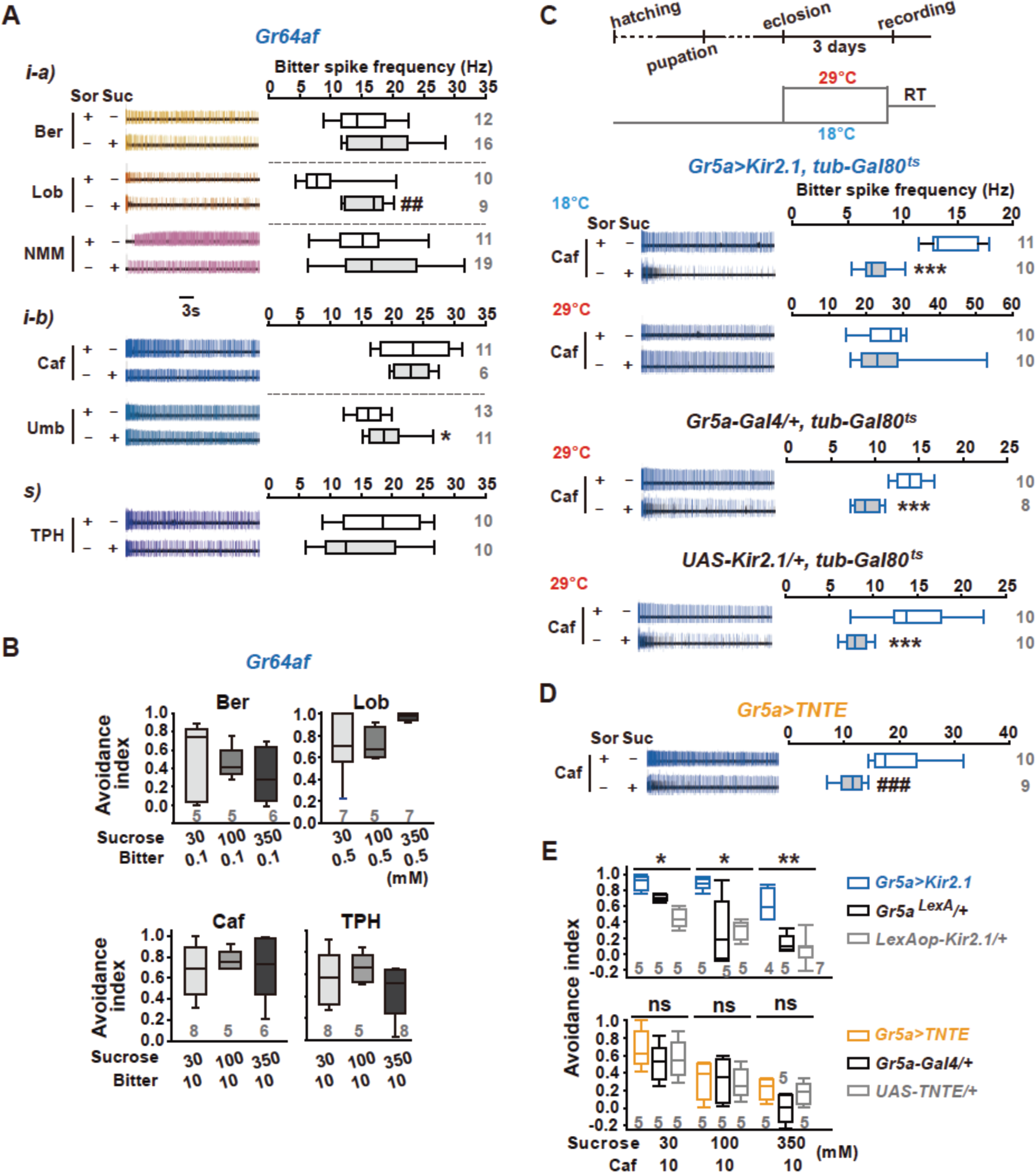
Excitation of sGRNs is required for sucrose-induced bGRN inhibition. (**A**) Sucrose-induced bGRN suppression in *Gr64af* mutant flies. Typical recordings (left) and box plots (right) are shown. The same chemical concentrations used in Figure 1. (**B**) Sucrose did not suppress feeding avoidance of bitters in *Gr64af* flies. (**C**) Silencing of sGRNs abolished sucrose-dependent inhibition of bGRN caffeine responses. *Gr5a-Gal4* is expressed in sGRNs. (Top) Experimental scheme for *Gal80^ts^* inactivation. (Bottom left) Typical recording results at the indicated temperatures. (Bottom right) Box plots from the experiments on the left. (**D**) Blocking chemical synaptic transmission with tetanus toxin light chain (TNT) did not affect sucrose-induced bGRN inhibition. (**E**) Silencing sGRNs but not blocking transmission reduced sucrose-dependent inhibition of caffeine avoidance. *, **, ***: Student’s t- or Tukey’s test, p < 0.05, 0.01, 0.001, respectively. ##, ###: Mann-Whitney U test, p < 0.01, 0.001, respectively. Gray numbers indicate the number of experiments.

### Optogenetic excitation of sGRNs inhibits bGRNs

Bitter-induced inhibition of sGRNs in L-type bristles requires the odorant-binding protein OBP49a, which is thought to bind various bitter compounds ^21^. It is possible that an OBP binds sucrose and the complex of OBP/sucrose inhibits bGRNs to achieve the observed inhibition. To determine whether OBPs contribute to sweet-dependent bGRN inhibition, we used an optogenetic approach with red activatable channelrhodopsin (ReaChR) ^45^ in combination with extracellular recordings (Fig. 3A-D). As sucrose is not present in the experiment, successful bGRN inhibition by optogenetic stimulation of sGRN would exclude involvement of a sucrose-binding mechanism. Applying a series of ten 0.1-sec orange light pulses at intensities ranging from 1 to 50 mW/cm^2^ 10 sec after initial contact with N-methyl maleimide (NMM), we measured bGRN spiking frequencies for 0.9 sec following each light pulse. NMM is an irreversible activator of TRPA1, which is natively expressed in bGRNs. As such, NMM generated sustained spiking responses ^36^. Activation of sGRNs by light stimulations of as low as 5 mW/cm^2^ significantly decreased the bGRN firing induced by 2 mM NMM, but maximal inhibition was achieved with stimulations of 25 mW/cm^2^ (Fig. 3B). The duration of the bGRN inhibition showed a monotonic correlation with increasing illumination intensity and time (Fig. S4B,C). In addition, we obtained similar results with caffeine and theophylline, which activate bGRNs in other bristle sensilla (Fig. 3C,D). From these results, we can conclude that the lateral inhibition of bGRNs we observed with optogenetic excitation of sGRNs does not require OBPs, and that the inhibition can be both rapid and prolonged.

**Fig. 3.**
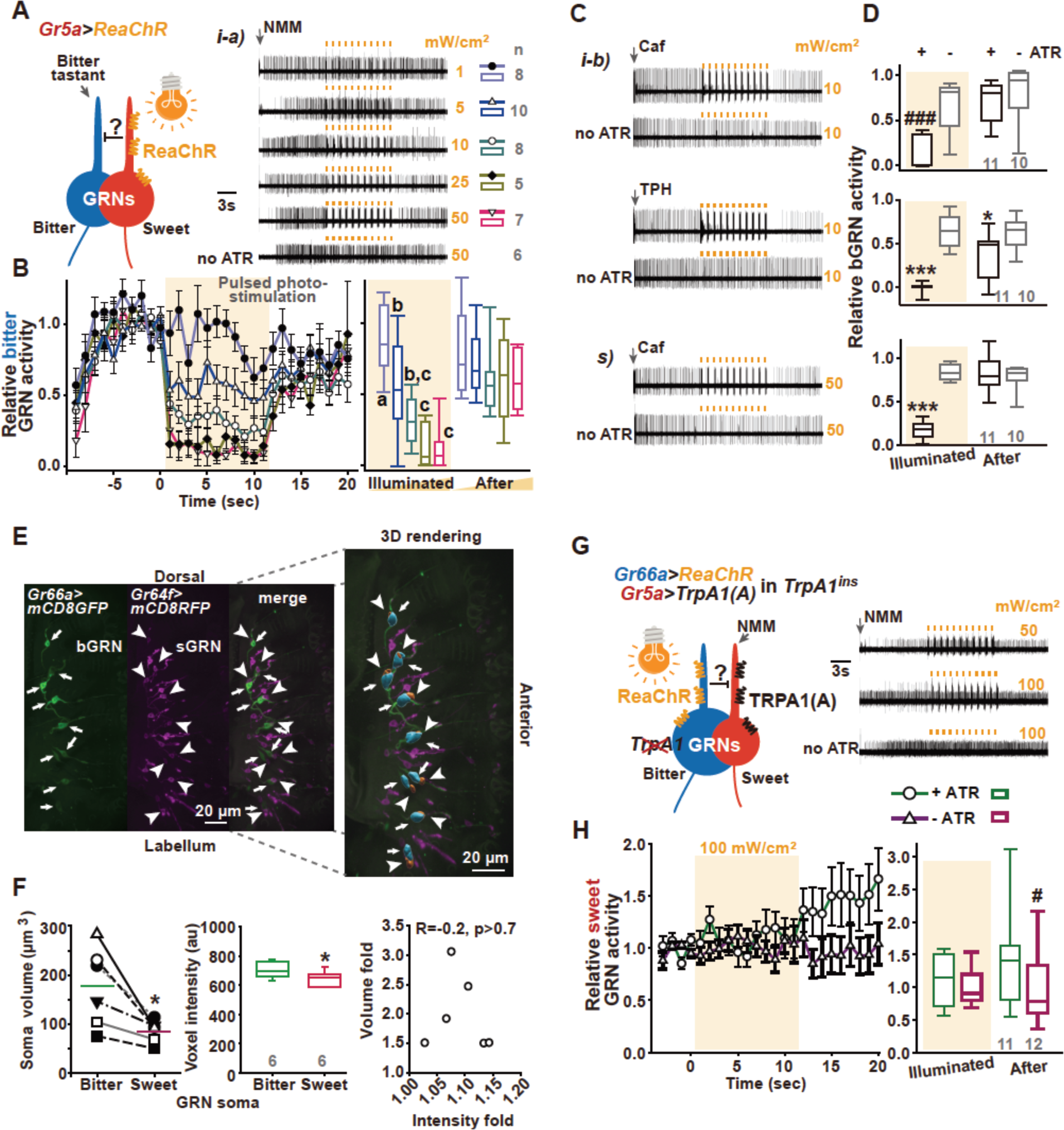
Optogenetically probed inhibition between GRNs is unilateral against the cell size gradient. (**A** and **B**) Optogenetic activation of sGRNs inhibits bGRNs activated by the TRPA1 agonist NMM (2 mM). Typical recordings showing irradiance dependence appear with a diagram illustrating the experimental design (A). ReaChR was expressed in sGRNs. 0.1-sec illuminations are indicated by small orange boxes. Average normalized spike frequencies of bGRNs for 0.9 seconds immediately after 0.1-sec illuminations at the indicated irradiance levels were plotted along the time axis (B). A translucent orange box indicates the time range over which light pulses were applied. Right, box plots showing average bGRN spiking frequencies during and after irradiation. (**C**) Typical optogenetic recordings following stimulation by the indicated bitters (Caf, 2 mM; TPH, 5 mM). (**D**) Box plots summarizing normalized data from (C). (**E**) bGRN soma volume was larger than that of its neighboring sGRN. Visualization of each GRN was achieved with the indicated membrane-tethered fluorescent proteins. 3D renderings for morphometric analyses were performed using Imaris 9.6.1. Arrows: bGRN soma. Arrowheads: sGRN soma. (**F**) Paired comparisons of soma volumes for juxtaposed GRNs (left). *: Paired t-tests, p < 0.05. bGRN and sGRN soma pairs with voxel intensities < 10% different were used for the paired analyses from the left (middle). Unpaired t-tests, p < 0.05. Volume and voxel intensity differences showed little correlation (right). (**G**) No bGRN-dependent sGRN inhibition was observed. Experimental design with TRPA1-expressing sGRNs and ReaChR-expressing bGRNs in *TrpA1*-deficient mutants (left). Typical recording results (right, NMM, 10 mM). (**H**) Average normalized spike frequencies of sGRNs for 0.9 second immediately after a 0.1-sec illumination at the indicated level of irradiance are plotted along the time axis. Letters indicate statistically different data groups, Dunn’s, p < 0.05 (B). *, ***: Student’s t-test, p < 0.05, 0.001, respectively (D). #, ###: Mann-Whitney U test, p < 0.05, 0.001, respectively (D,H). Gray numbers are n.

### The inhibiting sGRNs are smaller than the bGRNs they inhibit

Ruling out the involvement of synaptic transmission and OBPs, we began to consider ephaptic coupling as the mechanistic basis for this lateral inhibition, in part because ephaptic coupling has been reported in ORNs ^11,12^. Zhang et al. showed that spike amplitudes generally correlate with the relative cell size of the ORNs co-housed in a single sensillum, with the larger cells producing taller spikes. The larger ORNs readily suppress the smaller ORNs, resulting in an asymmetric yet bilateral ephaptic interaction ^12^. Intriguingly, our sensillar recordings of GRN activity seem to show that the sGRN produces lower amplitude spikes than the bGRN it inhibits (Fig. 1, Fig. S1A). To determine whether the larger cells produce the shorter spikes in the case of GRNs, we performed volumetric/morphometric analyses on 3D confocal images taken from flies with adjacent s- and bGRNs marked with different membrane-tethered fluorescent proteins (i.e., mCD8-GFP or mCD8-RFP) (Fig. 3E). We collected data only from GRN pairs with a voxel intensity difference of less than 10% (Fig. 3F, middle). The bGRN cell body volumes (mean ± SEM: 0.177 ± 0.033 mm^3^) averaged roughly two-fold larger than those of the sGRNs (0.086 ± 0.009 mm^3^, Fig. 3F, left). In other words, sGRN soma volume was like that of the large ORNs^12,46^. We did not see any correlation between the measured volume difference and the voxel intensities (Fig. 3F, right), excluding the possibility that our measurements were biased by differences in fluorescence intensity. Therefore, we conclude that, as with ORNs, spike heights are a reliable way of approximating the relative cell size of neighboring GRNs. This is presumably due, at least in part, to a similar distance of the recording electrode from the sites of origin for the action potentials in each neuron type. Thus, we confirmed that the smaller sGRNs exert lateral inhibition on the larger bGRNs.

### Lateral inhibition between sGRNs and bGRNs is unilateral against the cell size gradient

We next asked whether ephaptic inhibition between GRNs is bilateral, with the opposite direction (bGRN-dependent sGRN suppression) being more robust due to the size gradient, as it is in ORNs. To this end, we subjected sGRNs to lateral inhibition by optogenetically activating bGRNs. We did this while also ectopically expressing TRPA1 in the *TrpA1*-deficient mutant *TrpA1^ins^*^47^ to generate sustained sGRN spiking in response to NMM (Fig. 3G,H). Strikingly, we did not observe any inhibition of NMM-activated sGRNs induced by optogenetic excitation of bGRNs even at the highest level of irradiance (100 mW/cm^2^). We also confirmed this result with sucrose-activated sGRNs (Fig. 4C,D), indicating that the ephaptic interactions between GRNs are unilateral, with the smaller cell exerting the ephaptic effect.

**Fig. 4.**
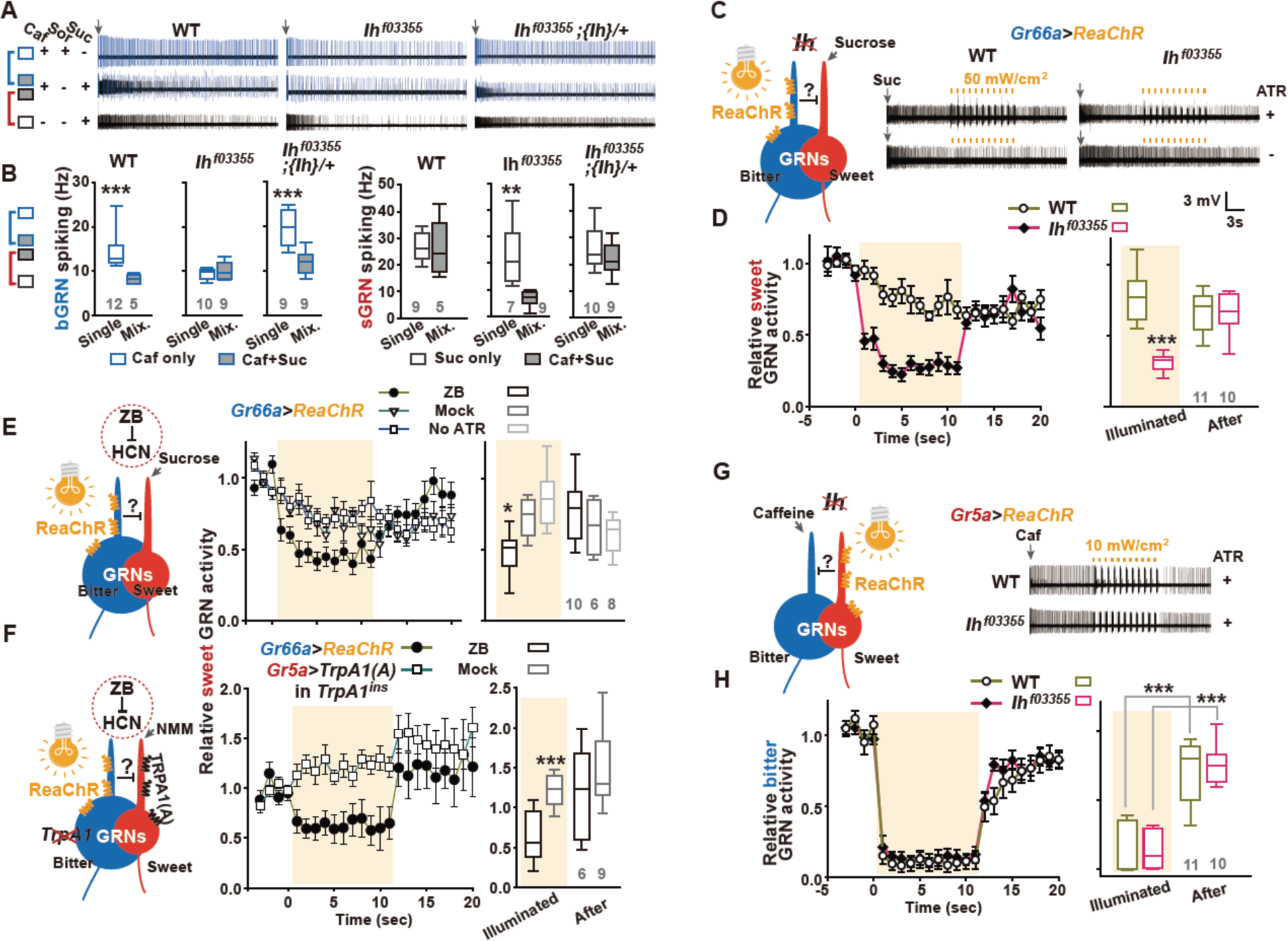
HCN loss-of-function makes the normally unilateral inhibition bilateral. (**A** and **B**) Lateral inhibition in the *Ih* mutant *Ih^f^*^03355^ is reversed. Typical recordings from the indicated genotypes (A). Box plots of average spike frequencies for the first 5 sec. *{Ih}*: a genomic fragment containing the entire *Ih* locus. Sor and Suc, 50 mM; Caf, 2 mM. (**C** and **D**) Optogenetic recordings reveal that bGRN-dependent sGRN inhibition is possible in *Ih^f^*^03355^. Suc, 10 mM. (**E** and **F**) The chemical inhibitor of HCN zatebradine (ZB) reversed the lateral inhibition in WT under optogenetic experimental settings. Suc, 10 mM; NMM, 10 mM. (**G** and **H**) sGRN-dependent bGRN inhibition was readily observed in *Ih^f^*^03355^, indicating bilateral inhibition in the mutant. Caf, 2 mM. *, **, ***: Student’s t-test between single and mixed stimulations, p < 0.05, 0.01, 0.001, respectively.

### Hyperpolarization-activated cyclic nucleotide-gated channel (HCN), encoded by Ih, is required for unilateral ephaptic inhibition

In searching for the mechanism underlying unilateral inhibition between GRNs, we hypothesized that the hyperpolarization imposed on sGRNs by bGRN activation could be negated by a hyperpolarization-induced mechanism that counteracts it. This would manifest in a lack of ephaptic inhibition of sGRNs by bGRNs, leaving only the inhibition of bGRNs by sGRNs. We reasoned that HCNs may provide such a mechanism because they depolarize the membrane by conducting Na^+^ and K^+^ upon hyperpolarization. This would potentially relieve sGRNs from any hyperpolarizing inhibition. Although the human HCN family comprises four members ^48^, the *Drosophila* genome encodes only one HCN orthologue from the *Ih* locus that shares biophysical properties with mammalian HCNs in heterologous expression systems ^49^. *Ih^f^*^03355^ is a strong loss-of-function allele in which the *Ih* gene is disrupted by a transposon insertion ^50,51^. Interestingly, in *Ih^f^*^03355^, unlike WT, sucrose did not reduce bGRN responses to caffeine (Fig. 4A,B, Fig. S5A). Furthermore, the direction of lateral inhibition between the GRNs appeared to be reversed, with *Ih^f^*^03355^ showing reduced sucrose-induced sGRN spiking in the presence of caffeine (Fig. 4B, right). Introduction of a genomic fragment containing the *Ih* locus ({*Ih*}) rescued these impairments, confirming that HCN is required for sweet-dependent bGRN inhibition (Fig. 4A,B). Corroborating this result, we found that, in contrast to WT, optogenetic excitation of *Ih^f^*^03355^ bGRNs expressing ReaChR under the control of *Gr66a-LexA* ^52^ inhibited sucrose-stimulated sGRNs (Fig. 4C,D) regardless of bristle type (Fig. S5B). We next attempted to suppress HCN activity using the well-known chemical inhibitor zatebradine (ZB) ^53^. Bristle sensilla of interest were pre-exposed to either water (mock) or 10 mM ZB for 2 min, immediately followed by optogenetic extracellular recording. In flies exposed to ZB, optogenetic activation of bGRNs suppressed sucrose- and NMM-induced sGRN spiking, the latter of which was accomplished via TRPA1 expressed in *TrpA1-*deficient sGRNs under the control of *Gr5a-Gal4* (*32*) (Fig. 4E,F). This pharmacological result suggests the defect we observed in *Ih^f^*^03355^ was neither developmental nor remotely inflicted. The relative GRN spike heights in *Ih^f^*^03355^ flies were like those of WT, suggesting that the size gradient between the GRNs persisted in *Ih^f^*^03355^ (Fig. 4A). Next, we asked whether the bGRN suppression induced by optogenetic sGRN activation was affected in *Ih^f^*^03355^. Interestingly, sGRN excitation via ReaChR readily reduced caffeine-induced bGRN responses in *Ih^f^*^03355^ (Fig. 4G,H, Fig. S5C), indicating that ephaptic coupling between GRNs is bilateral in the absence of functional *Ih*. The apparent reversal of the unilateral inhibition with chemically excited GRNs in Fig. 4A,B suggests that the balance of the GRN interaction in the mutants shifted toward bitterness dominance.

### Ih acts in sGRNs to mediate unilateral inhibition

Our hypothesis is based on the putative expression of HCN in sGRNs. We performed optogenetic extracellular recordings using flies with GRN-specific RNAi knockdown of *Ih*. Driving *Ih* RNAi in sGRNs using *Gr64f-Gal4*^54^ allowed similar bilateral inhibition (Fig. 5A) to what we observed with *Ih^f^*^03355^ flies (Fig. 4C,D), indicating it is in sGRNs that *Ih* is required for unilateral ephaptic inhibition. Driving *Ih* RNAi in bGRNs with *Gr89a-Gal4*^32^ did not produce any difference compared to controls (Fig. 5B). In *Ih^MI^*^03196^*^-TG^*^4.0^ (*Ih-TG4.0*) flies, the protein trap *T2A-Gal4* is inserted into one of the *Ih* introns, resulting in reliable spatiotemporal reporting of *Ih* expression^55,56^. The expression of the reporter *Ih-TG4.0* was widespread in the labellum, when probed with *UAS-mCD8::GFP* (Fig. 5C,D). To investigate in which GRNs the expression of *Ih-TG4.0* lies, if so, the GFP images were superimposed with those of CD4::Tdtomato driven by either *Gr64f-LexA* (sGRNs, Fig. 5C) or *Gr66a-LexA* (bGRNs, Fig. 5D). Most cells labeled by *Gr64f-*driven Tdtomato overlapped with *Ih-TG4.0-*positive cells (Fig. 5C, Movie S1). On the other hand, bGRNs labeled by *Gr66a-LexA* appeared to colocalize only partially with GFP when the confocal stacks were examined image by image (Fig. 5D, Movie S2). The partial colocalization could be artifactual due to the widespread expression of *Ih-TG4.0* in both neurons and non-neuronal cells.

**Fig. 5.**
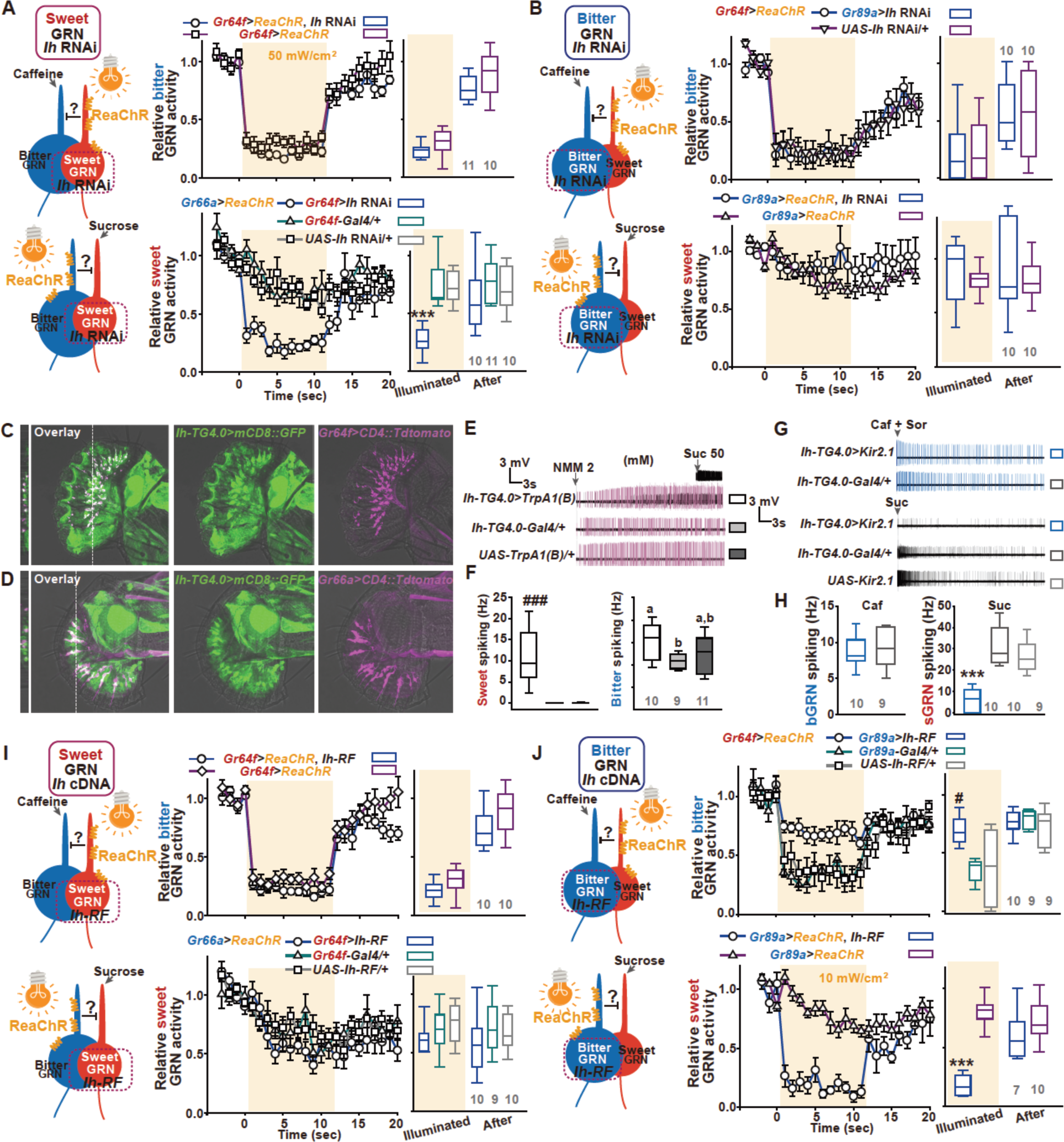
HCN expressed in sGRNs is required for unilateral ephaptic inhibition. (**A** and **B**) *Ih* RNAi in sGRNs made the GRN interaction bilateral. See the accompanying diagrams for each experimental design (left). (**C** and **D)** mCD8::GFP expressed by the *Ih* reporter *Ih-TG4.0* is colocalized in sGRNs but much less so in bGRNs expressing CD4::Tdtomato by indicated promoter lines. See Movies S1,2 for each confocal plane in the merged images. **(E-H**) *Ih-TG4.0* drives expression more highly in sGRNs than bGRNs. When the reporter expresses TRPA1(B), we found sGRNs generate small spikes in response to NMM in addition to the large spikes of the bGRNs (E and F). Kir2.1 expressed by the reporter dramatically reduced spiking in response to sucrose stimuli, but not to caffeine (G and H). (**I** and **J**) Gain-of-function experiments with the *Ih-RF* cDNA raised the possibility that HCN-expressing GRNs resist ephaptic inhibition. See the accompanying diagrams for each experimental design (left). ***: Unpaired t- or Tukey’s test, p < 0.001. #, ###: Dunn’s, p < 0.05, 0.001, respectively. Letters mark statistically different groups (D): Welch’s ANOVA, Games-Howell test, p < 0.05. NMM, 2 mM; Suc, 50 mM; Caf, 2 mM. Optogenetic actuation was done at the intensity of 50 mW/cm^2^ except for the last panel (J, bottom).

Therefore, we further examined the *Ih-TG4.0* expression in GRNs using extracellular recordings aided by molecular genetics. This functional approach provides a quantitative assessment of *Ih-TG4.0* expression in GRNs. In response to NMM, the i-type bristles of animals expressing *TrpA1(B)* under the control of *Ih-TG4.0* produced short spikes resembling sucrose-induced sGRN spikes (Fig. 5E) in addition to larger bGRN spikes resulting from native TRPA1 expression ^36^. This suggests *Ih* is expressed in sGRNs (Fig. 5E,F) because i-type bristles contain only two GRNs. Moreover, driving the expression of Kir2.1 with *Ih-TG4.0* produced a minimal reduction of the bGRN response to caffeine compared to the *Gal4*-only control and a dramatic reduction in sucrose-evoked firing (Fig. 5G,H). These data are consistent with a much higher expression of *Ih-TG4.0* in sGRNs than in bGRNs. According to the results of our GRN-specific *Ih* RNAi and reporter analyses, HCN expression in sGRNs is critical for counteracting the hyperpolarization putatively induced by bGRN excitation, leading to unilateral ephaptic inhibition.

### Ectopic expression of Ih in bGRNs confers resistance to sGRN excitation-dependent ephaptic inhibition

Since HCN expression in sGRNs is necessary for unilateral inhibition, we next examined the effect of HCN misexpression in bGRNs. After generating transgenic *UAS-Ih-RF* flies ^51^, we drove HCN expression in WT sGRNs using *Gr64f-Gal4.* The recordings of these *Ih-RF*-expressing sGRNs were indistinguishable from those of the control genotypes, indicating no effect on directionality (Fig. 5I). On the other hand, ectopic expression of *Ih-RF* in bGRNs using *Gr89a-Gal4* significantly attenuated sGRN excitation-dependent bGRN inhibition (Fig. 5J, top, Fig. S6). This was in contrast with the ephaptic bilaterality we observed in *Ih*-deficient GRNs (Fig. 3 and 5A), indicating that the *Ih* cDNA endows GRNs with resilience to ephaptic inhibition. In addition, *Ih* misexpression permitted sGRN inhibition upon optogenetic bGRN excitation (Fig. 5J, bottom), indicating that *Ih* can reverse the direction of ephaptic inhibition as well as help GRNs resist the inhibition. Thus, multiple lines of evidence have converged to indicate that differential HCN expression in GRNs determines the directionality of ephaptic inhibition.

### Chemosensory excitation of s- and bGRNs confirms the requirement of Ih expressed in sGRNs for unilateral interaction

We next sought to confirm our optogenetic findings by testing tastant-driven inhibition. Knock-down of *Ih* in sGRNs, but not bGRNs, appeared to reverse the direction of ephaptic inhibition (Fig. 6A,B), as we observed with *Ih^f^*^03355^ (Fig. 4A,B). Expression of *Ih-RF* in sGRNs, but not bGRNs, rescued *Ih^f^*^03355^ (Fig. 6C,D), confirming that HCN acts in sGRNs for the inhibition. Consistent with our optogenetic results with *Ih-RF* misexpression in WT bGRNs, we found chemically stimulated GRNs misexpressing *Ih-RF* also showed inverted lateral inhibition (Fig. 6E,F). Thus, our experiments with chemical stimulation were consistent with the optogenetic results above.

**Fig. 6.**
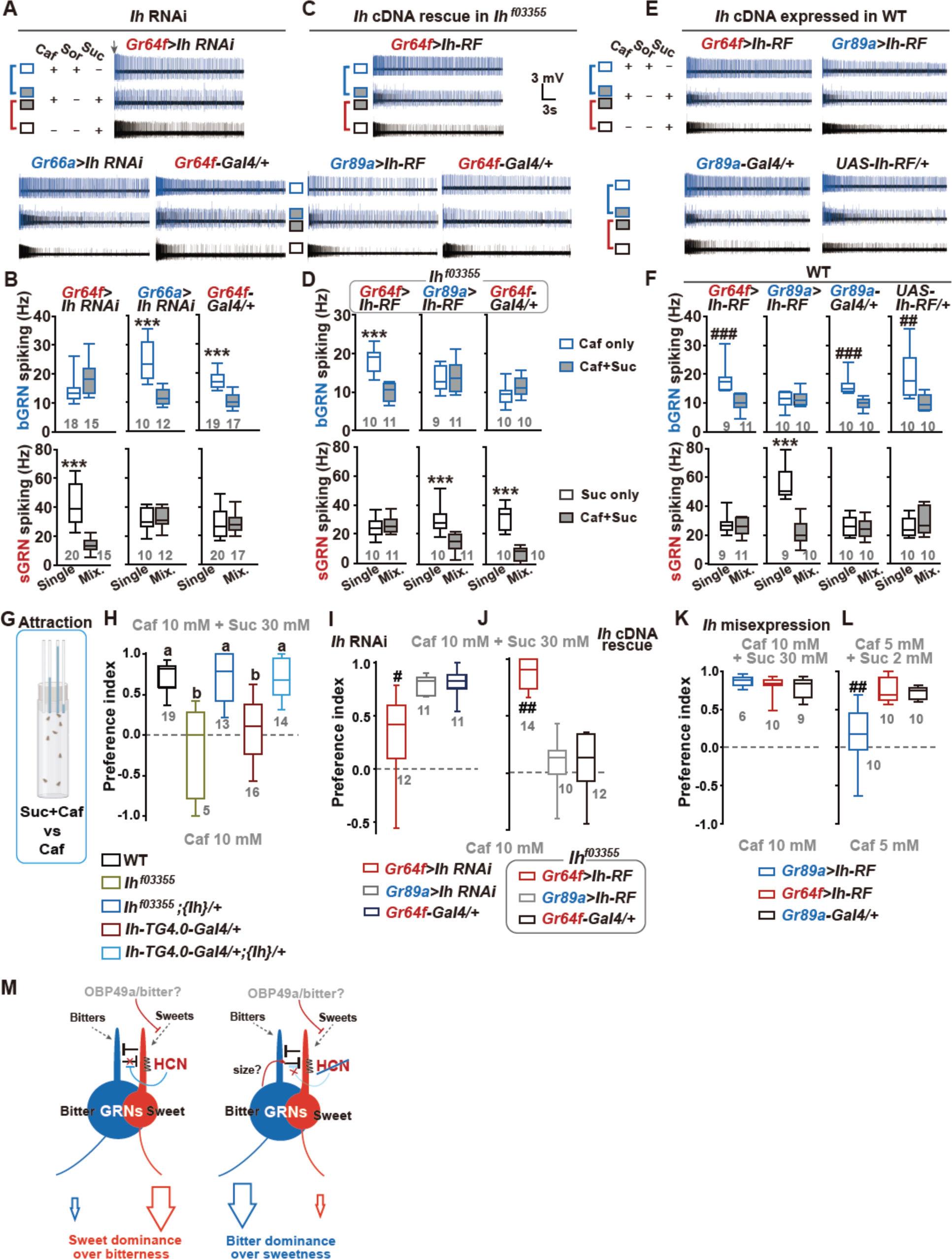
The unilateral ephaptic inhibition mediated by HCN expression in sGRNs allows animals to discern and ingest sweets buried in bitters. *Gr64f*- and *Gr89a-Gal4* were used to genetically manipulate sGRNs and bGRNs, respectively. (**A** and **B**) Expression of *Ih* RNAi in sGRNs reversed the lateral inhibition like *Ih^f^*^03355^ (***: unpaired t-test, p < 0.001, between tastant conditions). (**C** and **D**) Introduction of the *Ih-RF* cDNA in sGRNs (*Gr64f>Ih-RF)*, but not bGRNs (*Gr89a>Ih-RF*), in a *Ih^f^*^03355^ background restored sucrose-dependent bGRN inhibition (***, unpaired t-test, p < 0.001, between different tastants). (**E** and **F**) Misexpression of *Ih-RF* in bGRNs (*Gr89a>Ih-RF*) of WT reversed the direction of the lateral interaction (##, ###: Mann-Whitney U test, p < 0.01, 0.001, respectively. ***: Unpaired t-test, p < 0.001). Caf, 2 mM; Sor, 50 mM; Suc, 50 mM. (**G**) Schematic of the CAFE design for measuring preference index. (**H**) *Ih* deficiency impaired the ability of flies to ingest caffeine-laced sucrose. Letters indicate statistically distinct groups: Dunn’s, p < 0.05. (**I** and **J**) *Ih* in sGRNs is required for the ingestion of bitter-laced sucrose. (**K** and **L**) Misexpression of *Ih* in bGRNs of WT also impairs ingestion of a sucrose-bitter mixture. #, ##: Dunn’s, p < 0.05, 0.01, respectively. Gray numbers are n. (**M**) Diagrams illustrating the model of sweet/bitter interaction based on the results.

### Flies Ih-deficient in sGRNs failed to discern bitter-laced sucrose

We next examined the significance of the ephaptic inhibition in feeding using a CAFE assay designed to assess sucrose preference (Fig. 6G). To this end, we offered flies a choice between a sucrose (30 mM) solution with caffeine (10 mM) and a caffeine-only (10 mM) solution (Fig. 6H). Unlike CAFE illustrated in Figs 1 and 2, flies were examined in this way for their ability to choose sucrose-containing tubes while both sets of tubes contain the bitter caffeine. Remarkably, while WT were strongly attracted to the sucrose-caffeine solution, *Ih^f^*^03355^ and *Ih-TG4.0/+* were not (Fig. 6H). The defects introduced by both alleles were rescued by the *Ih-*containing genomic fragment {*Ih}*. We found *Ih* expression in sGRNs was necessary to achieve high preference indices, as both *Ih* RNAi knock down and *Ih* cDNA rescue in sGRNs, but not bGRNs, differed significantly from their respective controls (Fig. 6I,J). Although *Ih* misexpression in bGRNs reversed the lateral interaction between the GRNs (Fig. 5J and 6E,F), we observed a feeding defect when we adjusted the concentrations of caffeine and sucrose toward a balance of increased relative bitterness (Fig. 6K,L). This was probably due to a shift in the balance of the GRN interaction in a way that was distinct from that of *Ih-*deficient flies. These results suggest that the *Ih-* dependent GRN interaction is required for *Drosophila* to readily ingest sweets mixed with bitter chemicals.

## Discussion

Despite the growing number of reports on ephaptic coupling among neurons ^4,6,7,9,12,57^, little is known regarding the underlying molecular mechanisms. We report that neighboring sweet and bitter GRNs non-synaptically crosstalk in the form of ephaptic inhibition, and that the *Drosophila* HCN channel controls the directionality of ephaptic inhibition between these cells. The HCN current (known as I_h_) is well known to stabilize the membrane potential ^58^; HCN is activated upon and counters membrane hyperpolarization by conducting cations, while depolarization can decrease cation conductance through HCN standing at resting membrane potential, which thereby opposes the increasing membrane potential. Post-ephaptic neurons are hyperpolarized upon local field potential change induced by excitation of pre-ephaptic neurons, which is regarded as the electrophysiological mechanism for ephaptic inhibition^9,12,57^. HCN in sGRNs resists such ephaptic hyperpolarization, leaving only the direction of sGRN-dependent bGRN inhibition to be available (Fig. 6M). A hemichannel was previously proposed to be implicated in an ephaptic mechanism between cone cells and horizontal interneurons ^59^. The non-selective and constitutively conducting hemichannel in the post-synaptic membrane of horizontal cells appears to upregulate the presynaptic voltage-gated Ca^2+^ channel by subtly depolarizing cone cells across the synaptic cleft in a static manner. This suggests that the hemichannel mediates retrograde synaptic regulation, which contradicts the consensus that in ephaptic inhibition, electrical excitation of pre-ephaptic neurons dynamically inhibits adjacent neurons without the need for synapses. Thus, the hemichannel may have a clear role in regulating the chemical synaptic transmission, but does not pertain to ephaptic coupling.

Ephaptic inhibition in *Drosophila* ORNs is bilateral, but shows functional asymmetry between interacting ORNs^12^. This strategy of directionalization may lack the signal contrast that is achieved by HCN-dependent unilateral ephaptic inhibition, as significant fractions of inhibition can be mutually cancelled out between involved ORNs. The asymmetry in the interaction well correlated with the size gradient between the somas of the ORNs. In this regard, it is interesting that GRNs have opted for a genetic reprogramming of unilateral inhibition by means of the *Ih* gene, rather than having large sGRNs and small bGRNs. It is possible that a larger cell size is critical for bGRNs to retain high sensitivity detecting bitters, whereas bGRNs also need to be controlled by sGRNs. Neuronal process diameter generally shows a positive correlation with soma size ^60–63^. Thick dendrites may enable bGRNs to expose more bitter receptor molecules to the environment, lowering the bitter detection threshold. In addition, large axon diameters are often associated with higher rates of action potential propagation ^63,64^ and larger synaptic boutons ^65^, suggesting that soma size largely determines the rate of intraneuronal signal propagation and synaptic transmission robustness. Therefore, bGRNs may be larger because of the animal’s evolutionary need to sensitively detect and rapidly respond to the presence of potential toxins. Furthermore, flies appear to avoid bitter/sweet mixtures, when bitter levels are too high for the nutritional value represented by sweetness^18^. This is probably achieved by the bitter-binding protein OBP49a^21^. It will be interesting to see in future studies how ephaptic inhibition interacts with the OBP pathway, shaping sweet/bitter taste interaction for the feeding decision (Fig. 6M).

The evolutionary conservation of the HCN gene family with a pre-metazoan origin ^66,67^ suggests the implementation of HCN-dependent ephaptic regulation in neural information processing in the mammalian brain. Mammalian HCN channels have been implicated in many important brain functions such as learning and memory ^68,69^, sleep ^70,71^, and neural oscillations ^69,72–74^. As learning ^75,76^ and sleep ^77,78^ are known to correlate with specific frequencies of neural oscillations, HCN may contribute to diverse brain functions through its role in neural oscillations ^79^. Although intrinsic oscillations of single neurons by the interplay between HCN and voltage-gated Ca^2+^ channels were proposed to be a source of neural oscillations ^80^, HCN-dependent resilience to oscillating local field potential may contribute to shaping the waves, given that neural oscillations were reported at least in part to propagate through ephaptic coupling ^4,5,7,8,81^. Thus, whereas HCN channels and ephaptic coupling have been separately documented to be associated with neural oscillations, our findings of HCN-regulated ephaptic coupling in *Drosophila* gustatory organs imply the possibility of their collaboration for the mammalian brain functions.

Our work provides a foundation for future studies on the genetic regulation of ephaptic inhibition by demonstrating that the *Drosophila* gustatory system offers a simple and genetically tractable model of ephaptic inhibition. To date, the lack of genetic factors has hindered researchers from positively identifying ephaptic signaling. The ability of *Ih* to provide bGRNs with resilience to possible ephaptic inhibition proposes that *Ih* can be a genetic tool to experimentally discover ephaptic inhibition, as ephaptic inhibition is known to commonly involve hyperpolarization in the target neurons^3^. Thus, our work presents a novel genetic ephaptic program that meets the physiological needs of organisms, and aids future studies of ephaptic coupling by unraveling a key genetic factor in the directional control of ephaptic interaction. Furthermore, it is interesting to study in the future how genetically tuned ephaptic coupling cooperates with electrical and chemical synaptic transmission for the various information and behavioral coding of neural networks.

## Materials and methods

### Fly lines

Cantonized *w*^1118^ was used as wild type. *Gr5a^LexA^* and *Gr64f^LexA^*, *Gr64f-Gal4* were provided by Dr. Hubert Amrein^82,83^, while *Gr66a-LexA*^52^ by Dr. Kristin Scott. *Gr64af* is a gift from Dr. Seok Jun Moon ^34^. *TrpA1^ins^* and *UAS-TrpA1(B)* were previously described ^36^. *LexAop-CD4::Tdtomato* (Bloomington stock number #77139)*, UAS-mCD8::GFP* (#5137)*, LexAop-mCD8::GFP* (#32205)*, UAS-mCD8::RFP* (#32219)*, UAS-Rdl RNAi* (#89904, #31662), *UAS-GluClα* (#53356)*, UAS-ShakB RNAi* (#27291, #64087)*, UAS-Ih RNAi* (#58089), a duplicate of the *Ih* locus (denoted as *{Ih}* in the main text, #89744), *UAS-ReaChR* (#53749)*, LexAop-ReaChR* (#53747)), *Ih^f^*^03355^ (#85660) and *Ih-TG4.0* (#76162) were acquired from Bloomington Drosophila Stock Center (#stock number). The *UAS-Ih-RF* line was generated by Korea Drosophila Resource Center (http://kdrc.kr) by site-specific recombination into attP49b (3R), for which we cloned *Ih* cDNA through reverse transcription. The genotypes used in this study are listed in the following table.

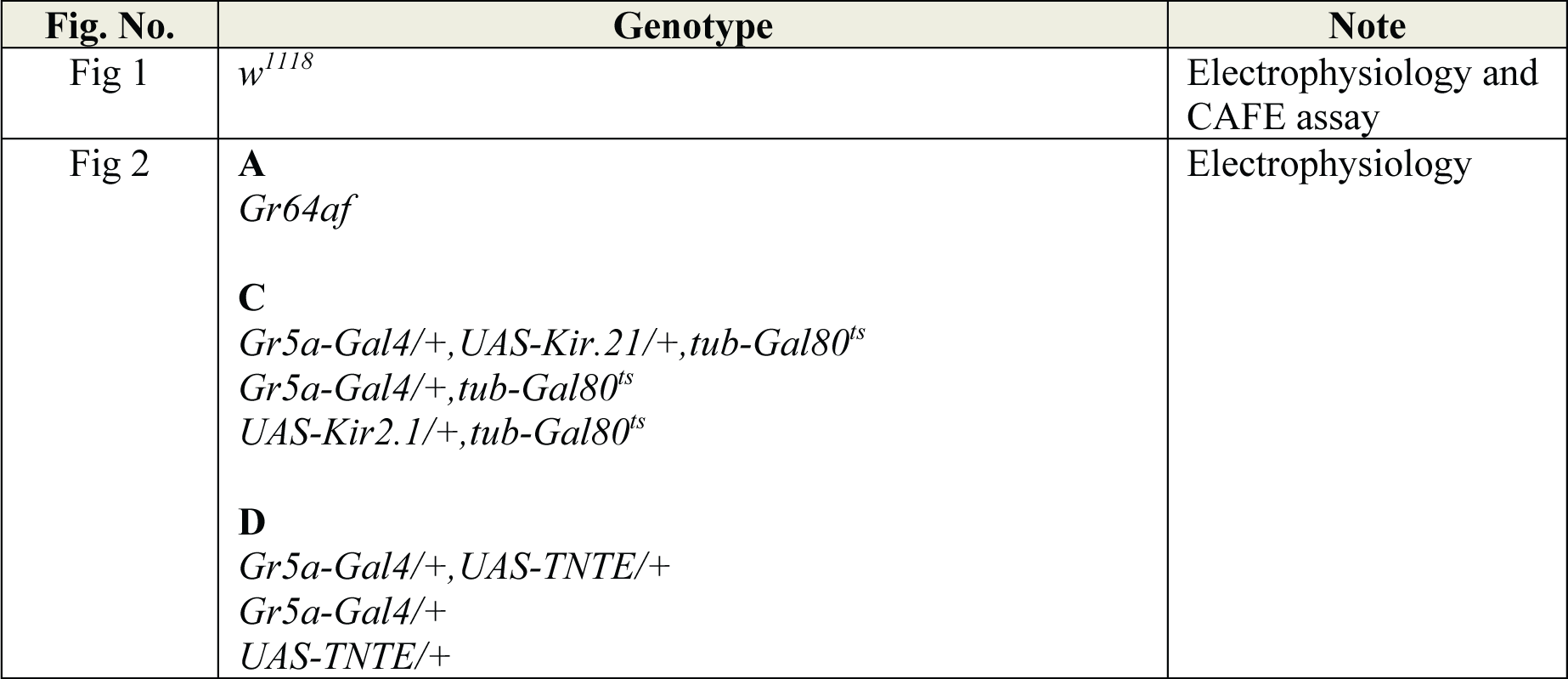
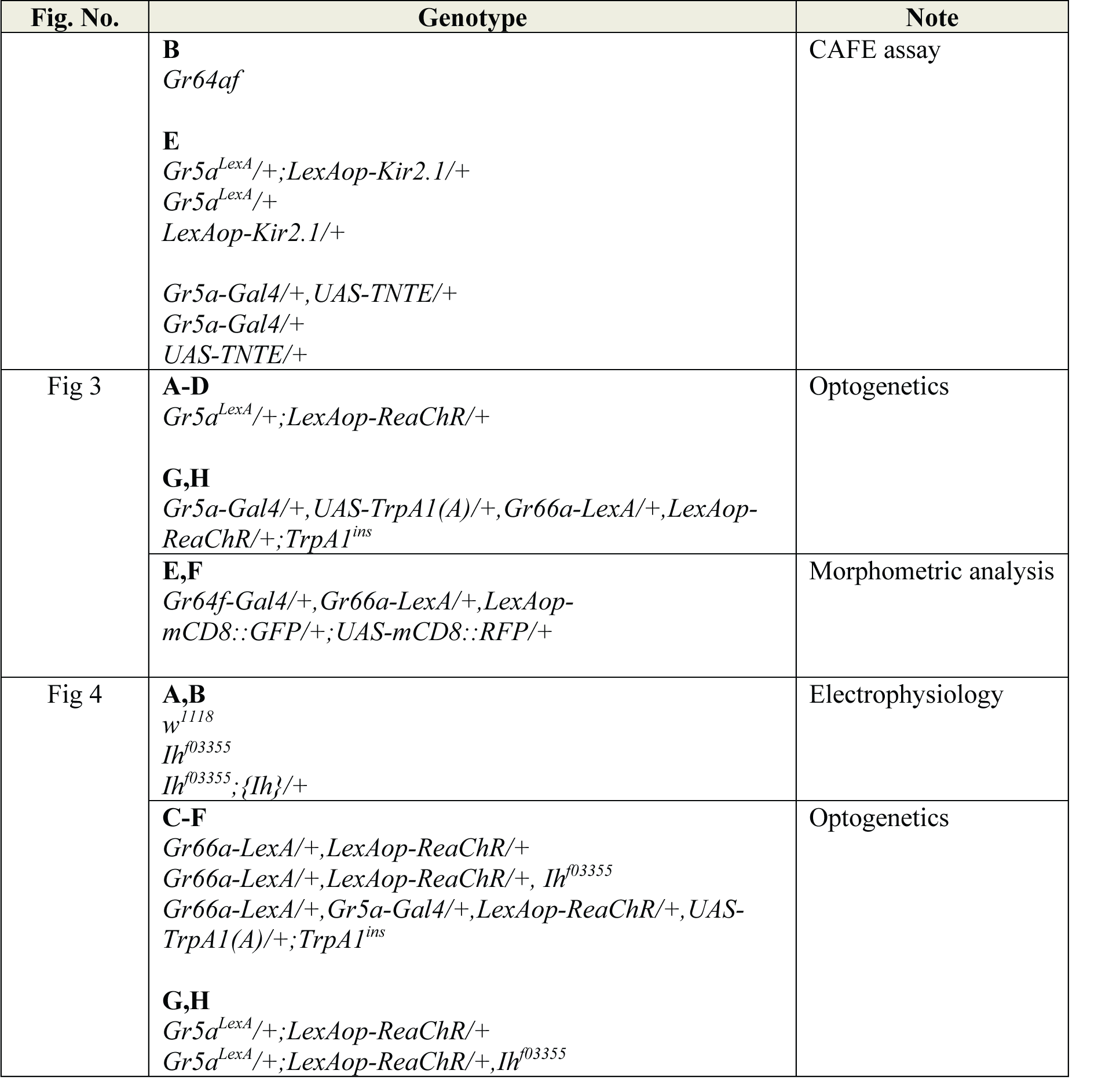
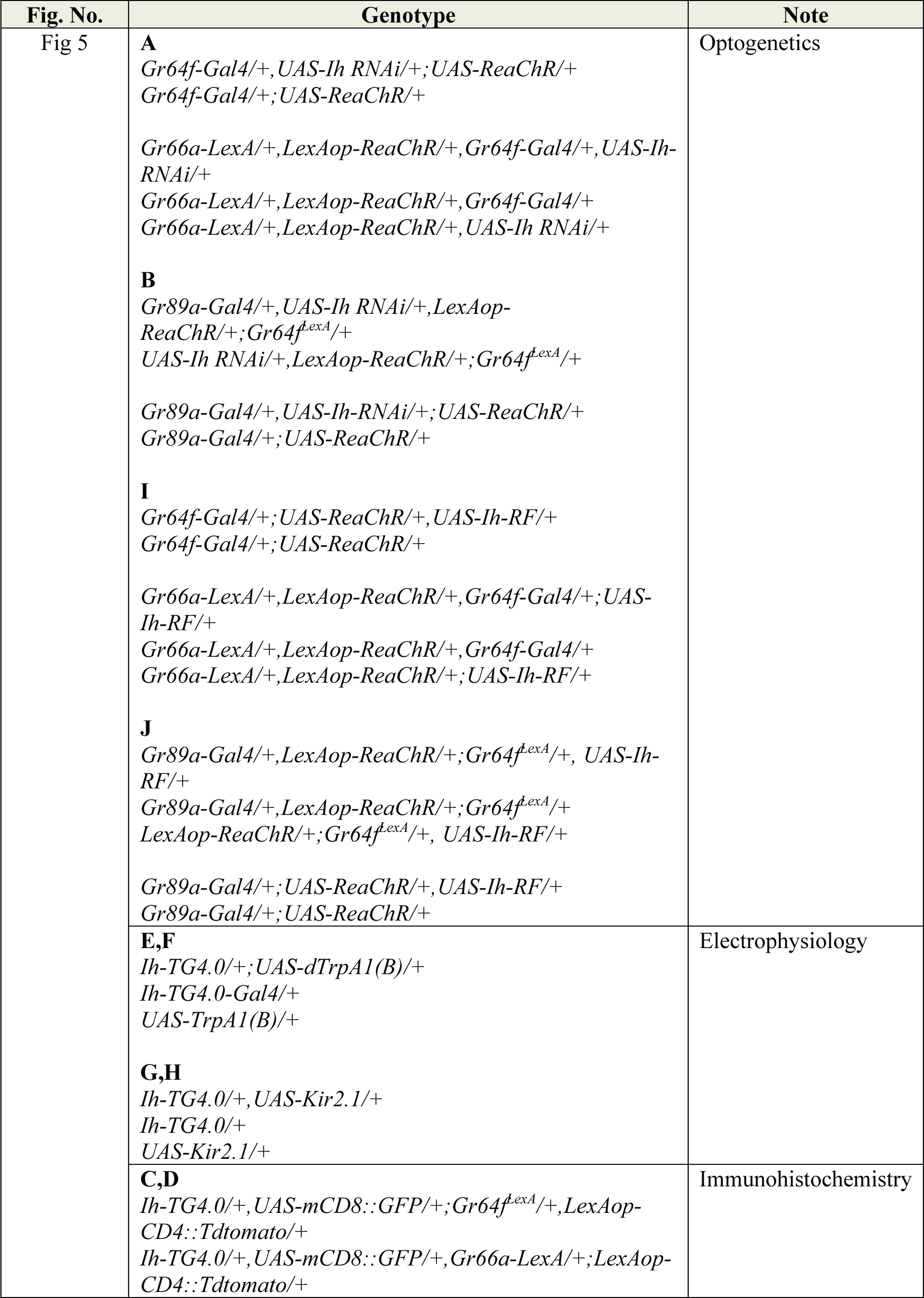
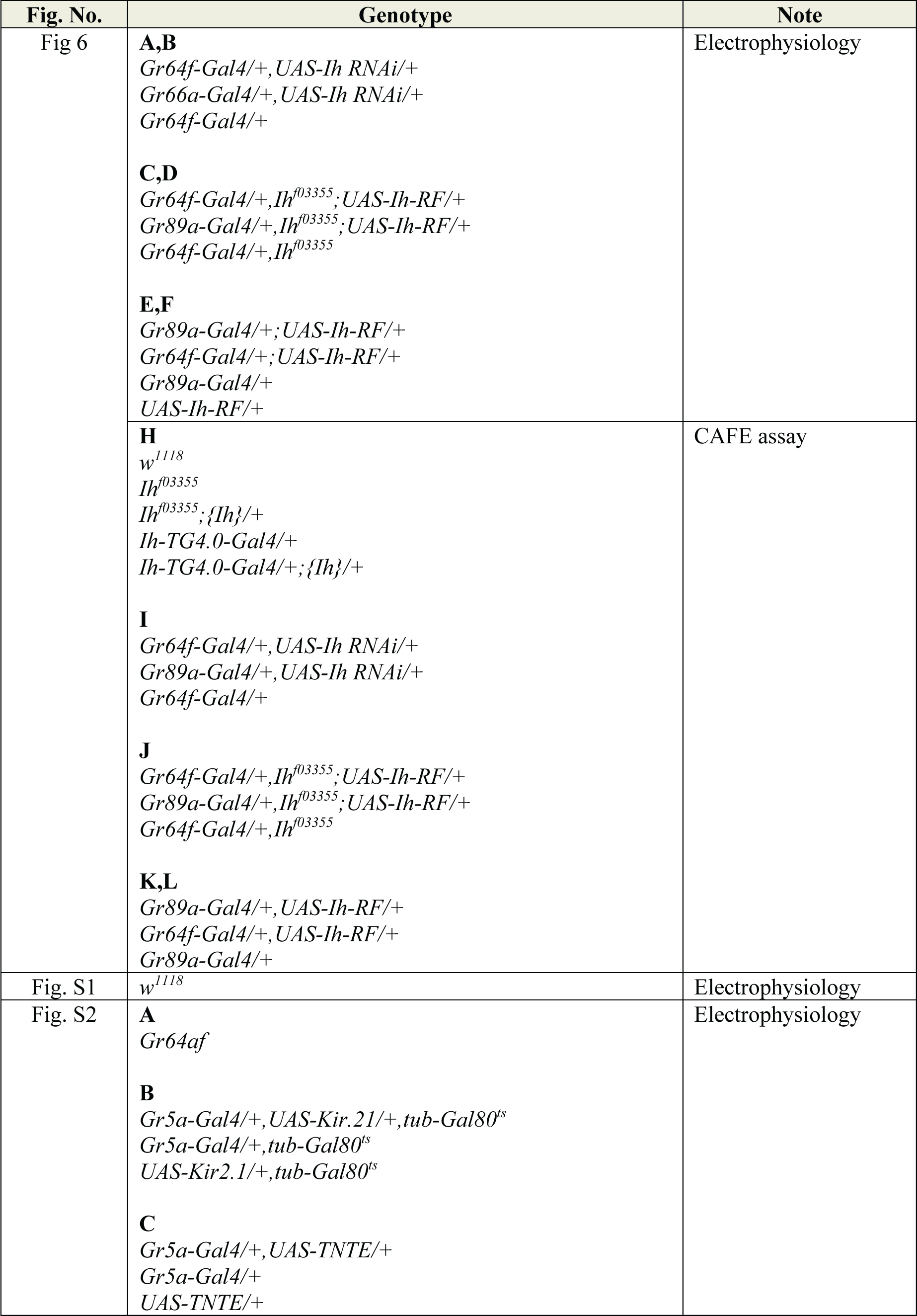
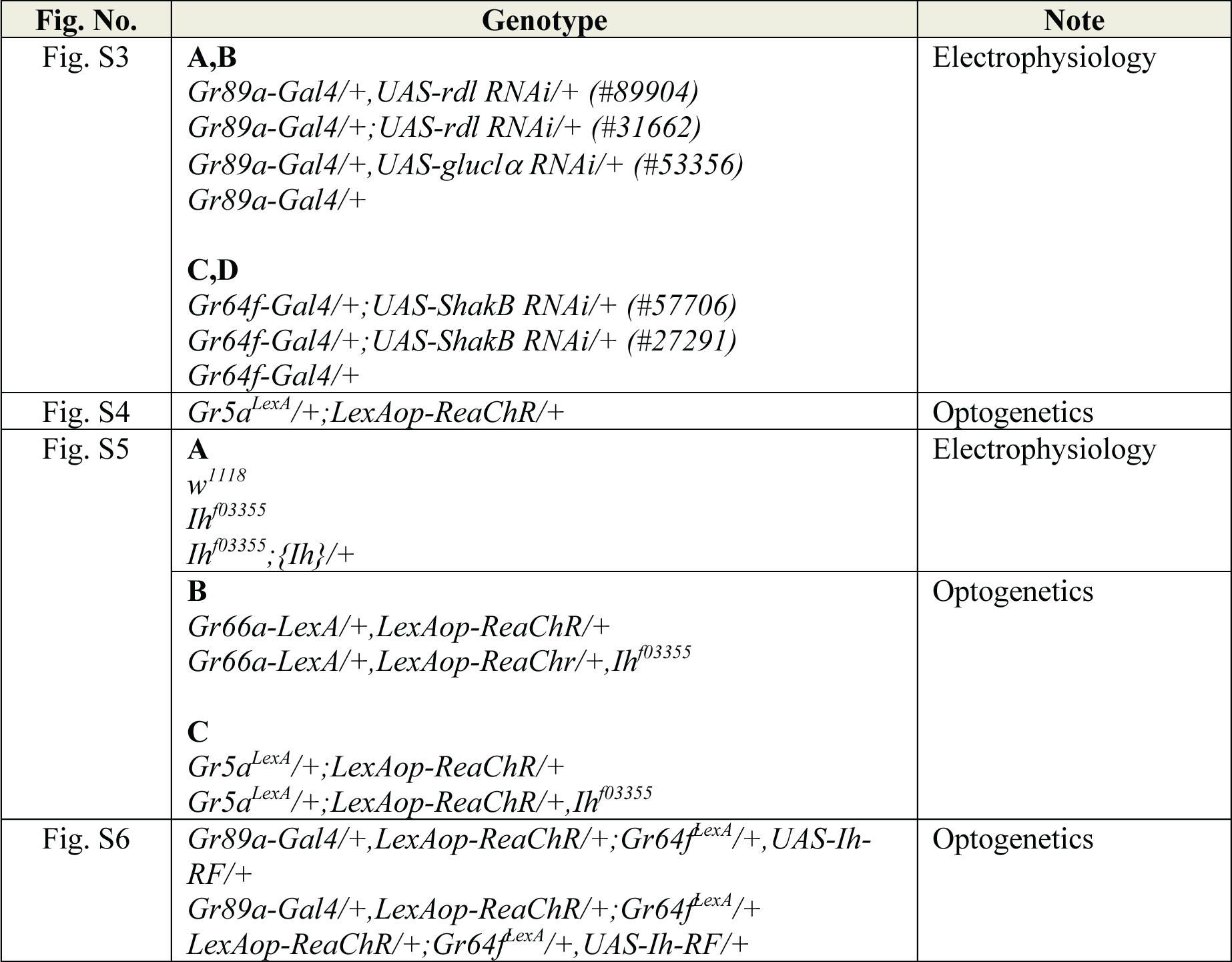

### Extracellular recordings and optogenetics

In vivo extracellular recordings were performed as detailed previously^84,85^. Each of the i-a, i-b, s-b type sensillum of 3-5 day-old flies were identified from the sensillum map described elsewhere^32^. The reference electrode was filled with HL3.1 solution^86^. The recording electrode contained tastants solubilized in the electrolyte tricholine citrate (TCC) at 2 (i-type) or 30 (s-type) mM. The concentrations of bitter chemicals were indicated in the corresponding figure legend. The information for acquisition of chemicals and fly lines is attached as a table at the end of Materials and Methods. For sGRN silencing experiments, *tub-gal80^ts^* was combined with *UAS-Kir2.1* and *Gr5a-Gal4* to avoid incomplete inhibition of the GAL4 activity. From hatching to eclosion, flies were incubated at the permissive 18°C, after which flies were removed to the incubator set at 29°C for 3 days. During the recording preparation, flies were handled at room temperature. The spiking frequency (Hz) was calculated every 5 seconds out of 30-sec long recordings and the first 5-sec frequencies were compared between genotypes or experimental conditions. Optogenetic GRN activation was performed at i-, and s-type bristles through the red-activatable channelrhodopsin (ReaChR) expressed in sGRNs or bGRNs. When a *Gal4* driver was in use for RNAi and cDNA misexpression, *LexA* lines were used to express ReaChR in the other GRNs: *Gr64f^LexA::VP^*^1683^ (Fig. 5B,5J) and *Gr66a-LexA* (Fig. 5A,5I). GRNs were illuminated by the Xenon lamp (Lambda LS, Sutter, CA, USA) via the orange light filter (ET 580/25X, Chroma). Illumination intensities of the orange light were calculated after measuring the power output with a S120VC photodiode sensor connected to a digital console PM100D (THORLABS, NJ, USA). Flies were fed with food containing 40 μM all-trans retinal (+ATR conditions) or regular food only with the solvent DMSO (“-ATR” or “no ATR” conditions) for 2 days after eclosion, during which flies were kept in the dark. The orange light was delivered as a series of 100-ms light pulses at 1 Hz for 10 sec. The spikes in 0.9 sec following illumination were acquired to estimate the inhibition, and plotted as a function of time (every second). For statistical comparison, the spiking frequencies for 0.9 sec immediately after 0.1 sec illumination were averaged, which constitutes a datum for box plots and were presented with the box plot from the frequencies obtained for 10 seconds after illumination (post-illumination). The spiking frequencies were normalized with the respect to the average of frequencies for 3 sec right before illumination. The signals picked up by the electrodes were amplified by Tasteprobe (Syntech) and digitized at a rate of 20 kb/sec by PowerLab with Labchart software (ADInstruments).

### Feeding behavior assay (CAFE assay)

Twenty of 3-5 days with 10 males and 10 females were starved for 20 hours in a vial containing water-soaked Kim’s wipes, and then given a choice of two sets of capillaries. To examine sucrose-dependent suppression of bitter avoidance, the first two capillaries were filled with sucrose and the second set of two capillaries was filled with sucrose and bitters at indicated concentrations (each concentration is indicated in Fig. 1F,G and all capillaries in a CAFE assay contained the same concentrations of sucrose). The choice was offered for 30 min. Evaporation of each tastant solution was estimated using CAFE setups without flies and subtracted from the consumed volume. Avoidance indices were calculated as the net fraction of consumed volume of sucrose-only solution, which resulted from subtracting the fraction of consumed volume of sucrose+bitter solution from the sucrose-only fraction. To examine the ability of flies to find bitter-laced sucrose, starved flies were subjected to CAFE for 1 hr with the first set of capillaries containing sucrose in addition to caffeine and the second set of capillaries containing only caffeine at indicated concentrations. Preference indices were quantified by calculating the net volume fraction of consumed caffeine+sucrose solution compared to that of caffeine only (Fig. 6G-L).

### Immunohistochemistry and morphometric analysis

To localize the expression of HCN in sGRNs and bGRNs, we utilized mCD8::GFP driven by *Ih-TG4.0* and CD4::Tdtomato driven by either *Gr64f^LexA::VP^*^16^ or *Gr66a-LexA*, respectively. The probosces of 7-10-day old flies were dissected in cold PBS, fixed with 4% paraformaldehyde for 25 minutes at RT, and promptly mounted for confocal imaging. Z-stack were taken at 1 μm intervals using a 40x objective lens (W CORR CS2, NA: 1.10, Leica, Germany) on the Stellaris8 microscope (Leica, Germany). The obtained z-stack images were used to create Movie S1 and S2 using imageJ. To measure soma volume of sGRNs and bGRNs, sGRNs and bGRNs were delineated by expressing mCD8::RFP and mCD8::GFP driven by *Gr64f-Gal4* and *Gr66a-LexA*, respectively (Fig. 4E). The probosces of 7-10 days flies were dissected in cold PBS and fixed for 4 hrs at 4°C in 4% paraformaldehyde. After permeabilized in PBST (0.5% Triton X100) for 1 hr, samples were incubated in PBS + 10% H_2_O_2_ for 1 hr and blocked in the blocking solution (90% PBST/10% goat serum) for 2 hrs. The mouse anti-GFP (2655S, Cell signaling technology, USA) was added to the blocking solution with the dilution of 1:1000 at 4℃ overnight. The secondary antibody conjugated with fluorescein diluted at 1:1000 in the blocking solution was applied and incubated at 4℃ overnight. The samples were mounted in Vectashield (Vector Laboratories, H-1400) on a slide under a coverslip. Confocal images were acquired along the z axis with 200 nm intervals with a 60x objective lens (MRD01605, NA: 1.40, Nikon, Japan) attached to Dragonfly 502W (Andor, UK) for 3D rendering. The obtained images were processed for assessment of soma volume using Imaris x64 9.6.1 (Oxford Instruments), in which algorithm was set to segment only a region of interest and to calculate the shortest distance after adding new surfaces. The threshold for voxel inclusion is based on absolute intensity and manually determined for optimal outlining of cell surface. Before obtaining volume, dendrites were excluded and only the soma was selected using the “Cut surface” function.

### Statistics

Statistical calculation was performed using Sigmaplot 14.5 (Systat Software). The sample sizes and the different biological replicates used for each experiment and the statistical tests are specified in the figure legend. Normal distribution and heteroskedasticity were assessed using Shapiro-Wilk and Brown-Forsythe tests, respectively, before parametric tests. When these tests were failed, non-parametric tests were performed. However, for some cases, heteroskedasticity with normality led us to perform Welch’s t-test (Sigmaplot 14) or ANOVA, the latter followed by Games-Howell test as a parametric analysis using the Excel spreadsheet available at www.biostathandbook.com/welchanova.xls. No outlier was excluded for statistical analyses.

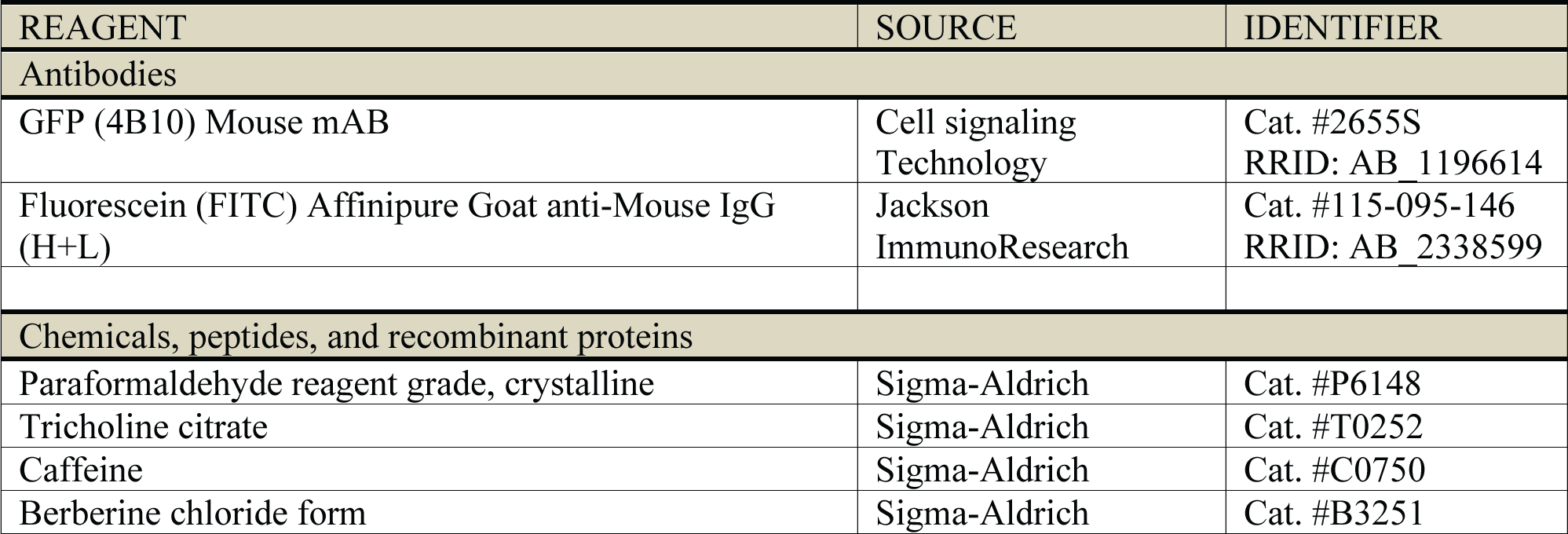
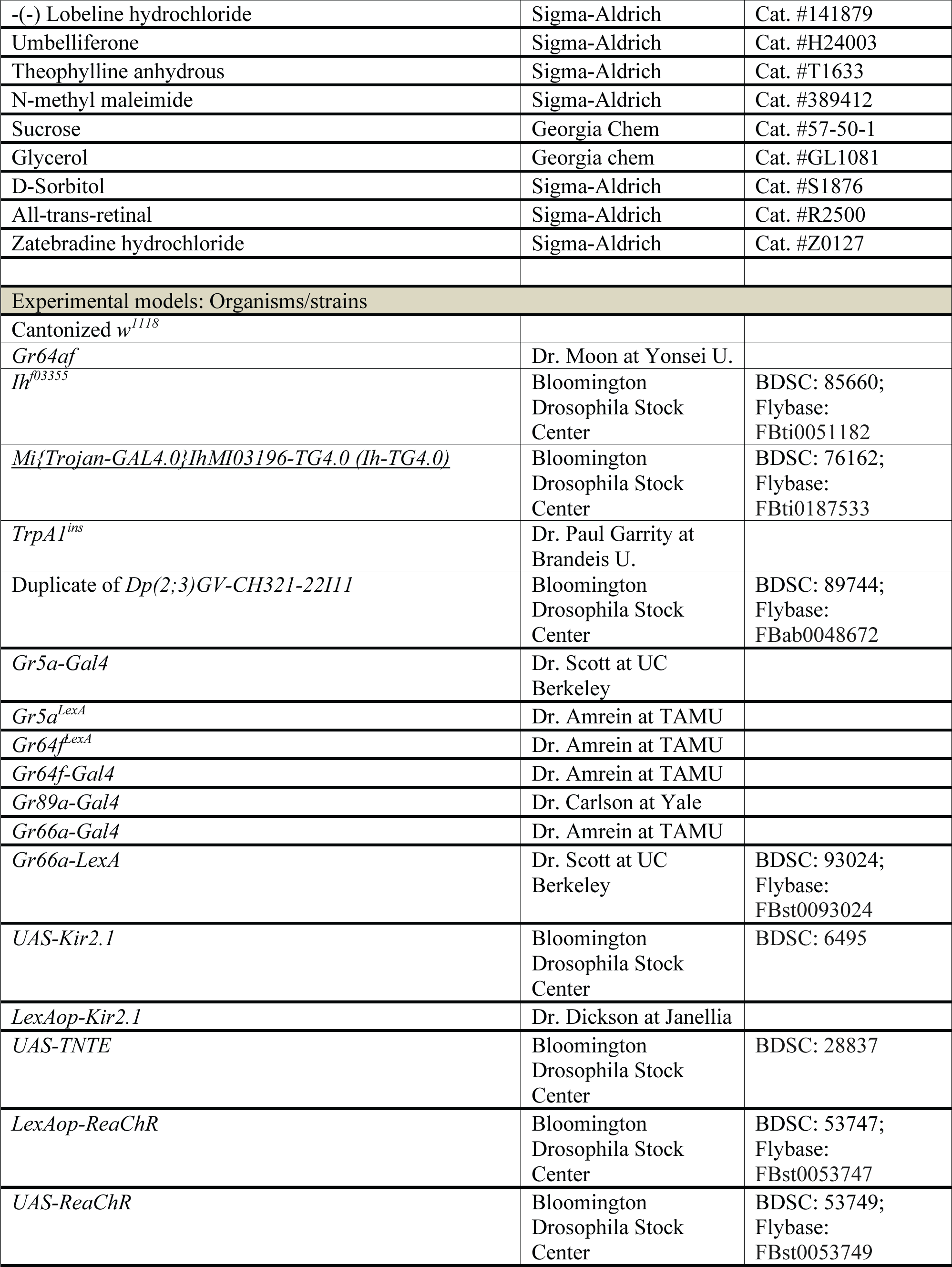
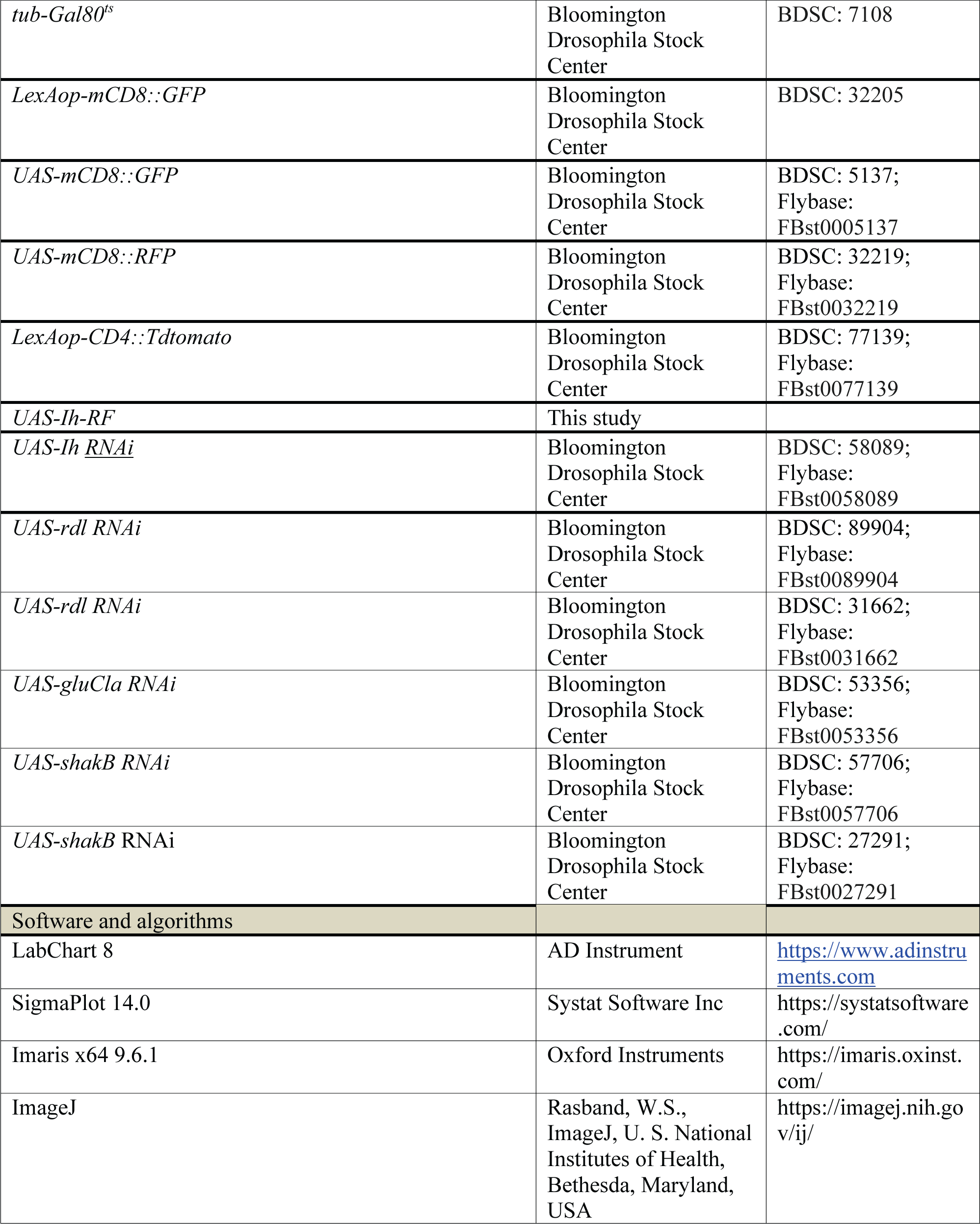

## Acknowledgments

We would like to thank Drs. Garrity, P., Lim, HH, Oh, WJ and Kim, G for helpful comments, and Drs. Amrein, H, Scott, K, Moon SJ, and KDRC/BDRC stock/resource centers for sharing fly lines as indicated in Methods and Materials

## Data and materials availability

All data are available in the main text or the supplementary materials.

## Author contributions

Conceptualization: MHL, KJK, KK

Methodology: MHL, KJK, SYK, TP, JYK

Investigation: MHL, KJK, KMJ

Visualization: MHL, KJK, SYK

Funding acquisition: KJK, KK

Project administration: JYK, KMJ, KJK, KK

Supervision: KJK, KK, JYK, KMJ

Writing: MHL, KJK, KK, JYK, KMJ, SYK, TP

