## Supplementary material for "Unilateral ephaptic program underlying sweetness dominance": supp figures 1-6

### Supp_Figure S1

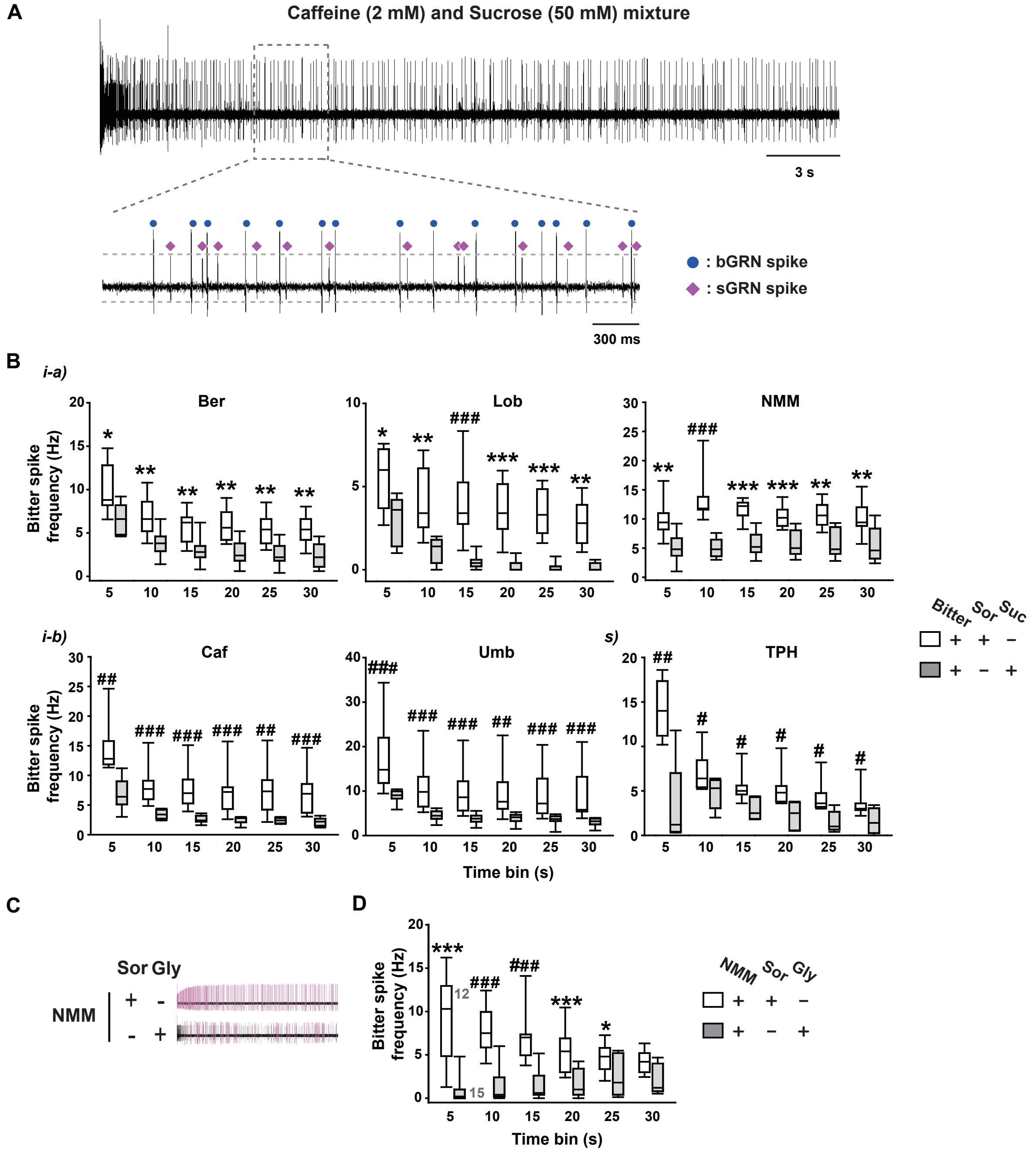


**Supplemental Fig. S1. sGRN-dependent suppression of bGRN activity.** (A) A representative recording of action potentials from WT i-type bristles stimulated by indicated sweet and bitter chemicals. The blue circles mark bGRN responses, while the magenta diamonds mark sGRN responses. The dashed lines indicate upper and lower boundaries of sGRN potential responses clearly distinct from those of bGRN responses. (B) Action potential frequency data for averaged bins every 5 sec for 30 sec, corresponding to the data in Figure 1B-D. 0.5 mM Ber, 0.5 mM Lob, 2 mM NMM, 2 mM Caf, 0.5 mM Umb, 5 mM TPH and 50 mM sucrose or sorbitol. * and #: $p<0.05$, ** and ##: $p<0.01$, *** and ###: $p<0.001$, Student's t- and Mann Whitney U test, respectively. (C) A representative trace for bGRN activity inhibited by glycerol that activates sGRNs. 112 mM Glycerol and sorbitol, 2 mM NMM. (D) Box plots of bitter spiking frequencies from (C) during indicated time bins. *: $p<0.05$, *** and ###: $p<0.001$, Student's t- and Mann Whitney U test, respectively.

### Supp_Figure S2

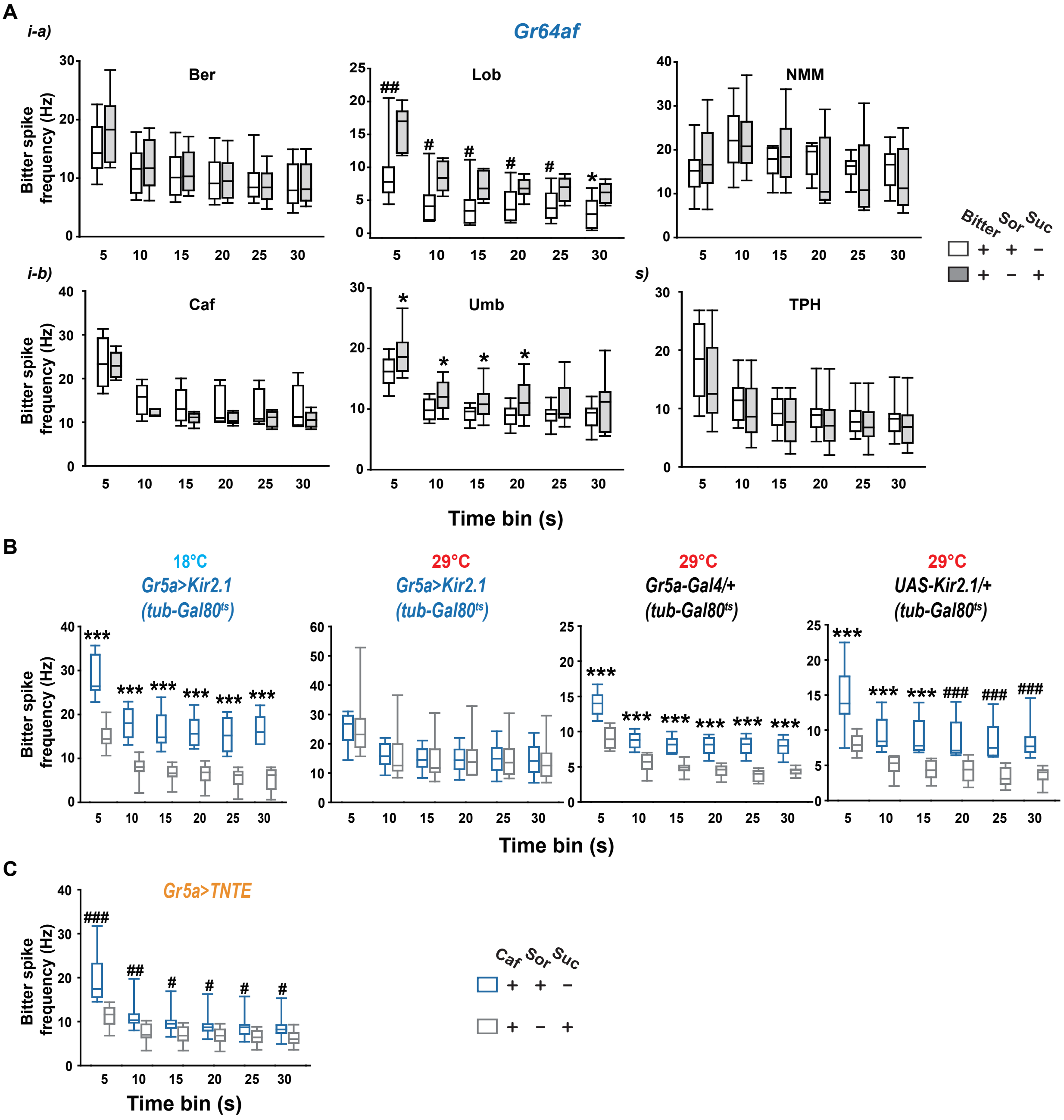


**Supplemental Fig. S2. Excitation, but not chemical synaptic inhibition, of sGRNs is required for inhibition of bitter GRN activity.** (A) Quantification of 5 sec bins from 30-sec recordings on i-type bristles in *Gr64af*, corresponding to the data in Fig. 2A. *: p<0.05, Student's t-test. #: p<0.05, ##: p<0.01, Mann-Whitney U test. (B) Quantification of 5 sec bins from 30-sec recordings on i-type bristles in *Gr5a>Kir2.1* (*Gr5a-Gal4/+,UAS-Kir2.1/+,tub-Gal80ts/tub-Gal80ts*) at indicated temperature condition. *** and ###: p<0.001, Student's t- and Mann-Whitney U test, respectively. Caffeine 2, Sor and Suc 50 mM. (C) Quantification of 5 sec bins from 30-sec recordings on i-type bristles in *Gr5a>TNTE*. #: p<0.05, ##: p<0.01, ###: p<0.001, Mann Whitney U test.

### Supp_Figure S3

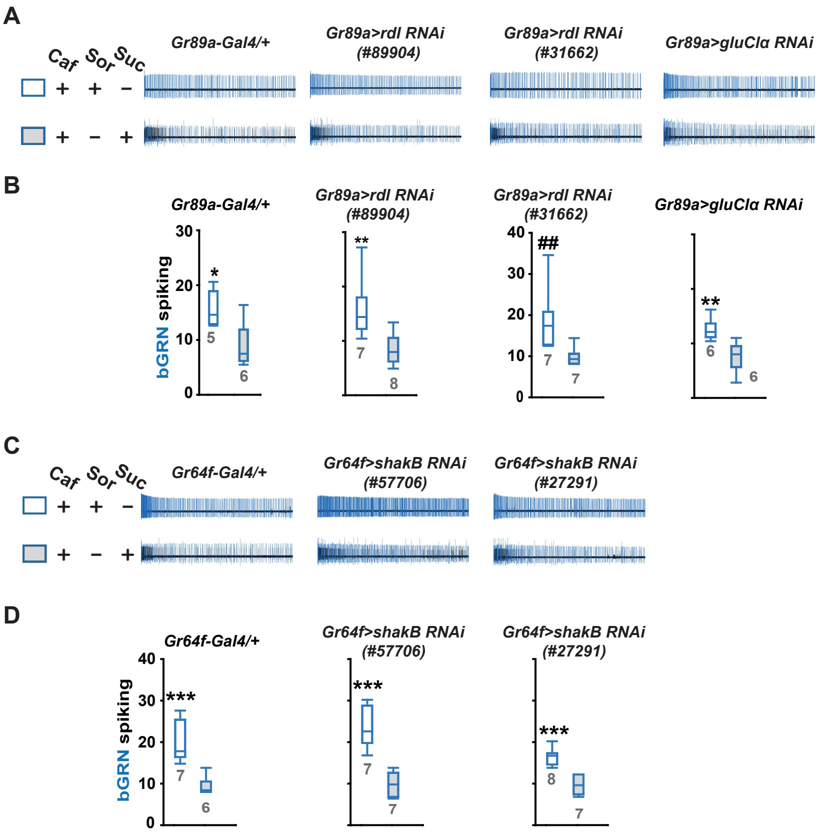


**Supplemental Fig. S3. RNAi knockdown of inhibitory postsynaptic receptor genes and a gap junction gene.** (A) Representative recording traces of RNAi knockdown in bGRNs of *rdl* encoding a GABA-activated chloride channel, and *gluClα* encoding a glutamate-gated chloride channel. (B) Quantification of a for caffeine-induced spiking frequency in the first 5-sec bin. *, **: Student's t-test. $p<0.05$, 0.01. ##: Mann-Whitney U test, $p<0.01$. (C) Representative recording traces of RNAi knockdown in sGRNs of *shakB* encoding a major neuronal innexin. (D) Quantification of (C) for caffeine-induced spiking frequency of the first 5 sec. ***: $p<0.001$, Student's t-test.

### Supp_Figure S4

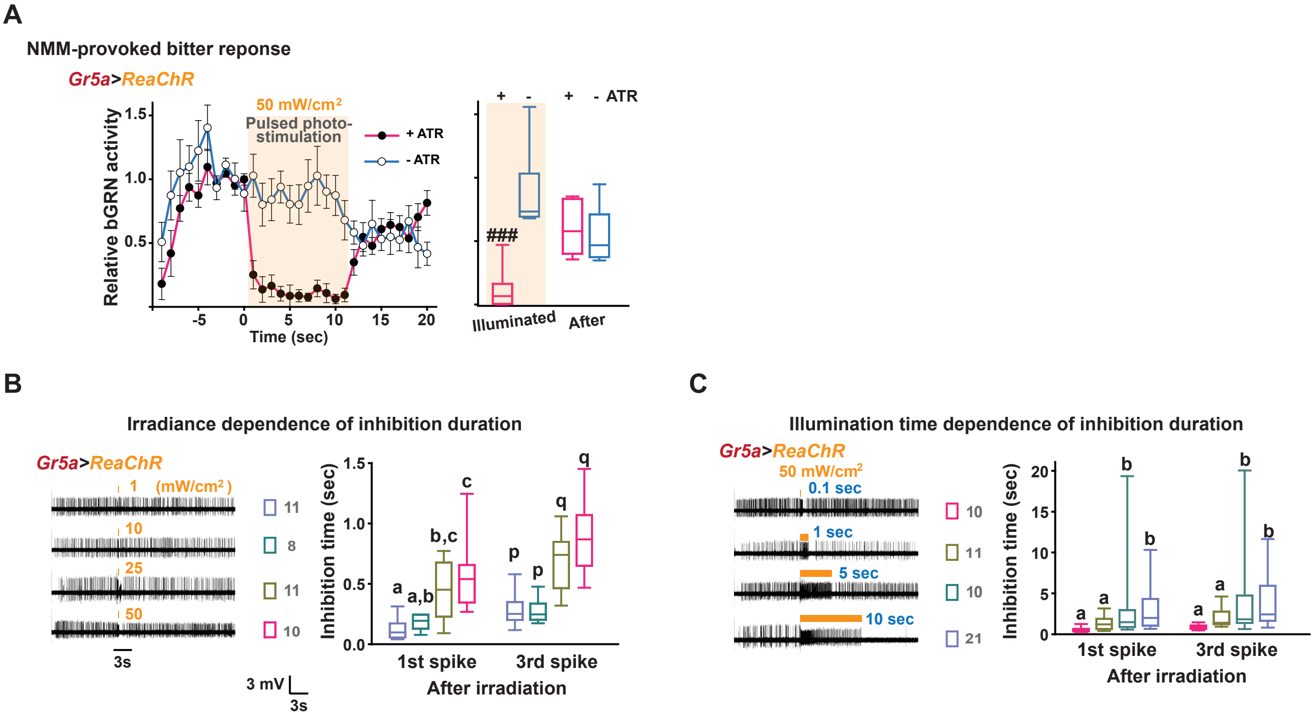


**Supplemental Fig. S4. Optogenetic characterization of lateral inhibition.** (A) Optogenetic inhibition of bGRNs is all-trans-retinal-dependent. Left: Spiking frequencies at each sec was plotted. Data with illumination is illustrated as an orange box. Right: quantification of indicated data during and after illumination. ###: $p<0.001$, Mann-Whitney U test. (B) Inhibition duration was measured as time duration until the first or third spikes of bGRNs after illumination. The irradiance was varied as indicated. Left, representative raw traces. Right, quantification of inhibition time with increasing light intensity. Letters indicate statistically distinct groups. a-c: Tukey's test, $p<0.05$. p and q: Dunn's test, $p<0.05$. Gray numbers are repeated times of the experiments. (C) Irradiation time dependence of inhibition with increasing illumination time. Left, representative recordings. Right, quantification from replicates of Left. Letters indicate statistically different groups: Tukey's test, $p<0.05$. Gray numbers are repeated times of the experiments.

### Supp_Figure S5

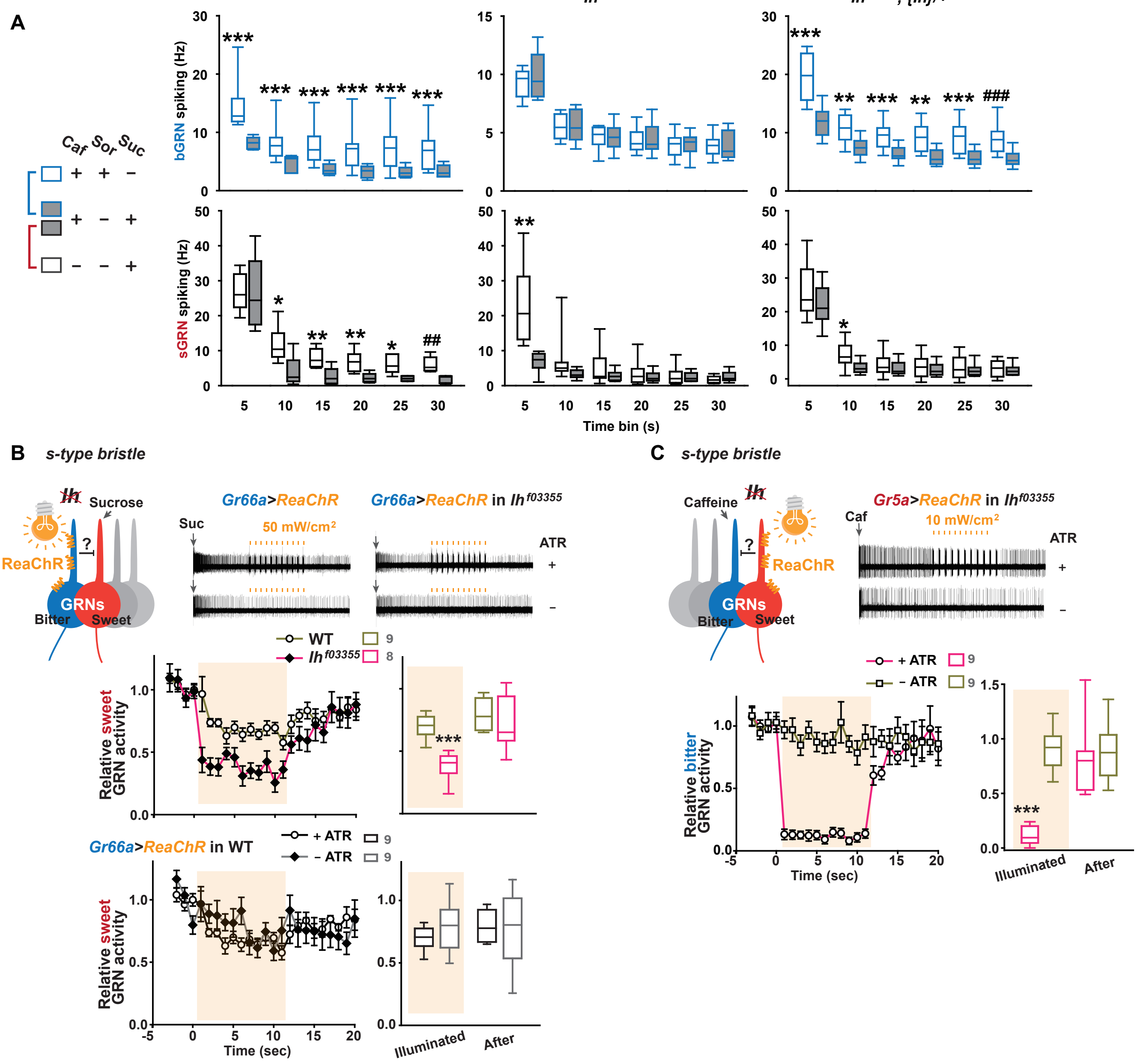


**Supplemental Fig. S5. *Ih* is required for unilateral inhibition between sGRNs and bGRNs.** (A) Box plots of the mean bGRNs or sGRNs spike for indicated time bins, corresponding to the data in Fig. 4A,B. *: p<0.05, Student's t-test, ** and ##: p<0.01, *** and ###: p<0.001, Student's t- and Mann Whitney U test, respectively. (B and C) Lateral inhibition between sGRNs and bGRNs in s-type bristles is also bilateral without functional *Ih*. ***: p<0.001, Student's t-test.

### Supp_Figure S6

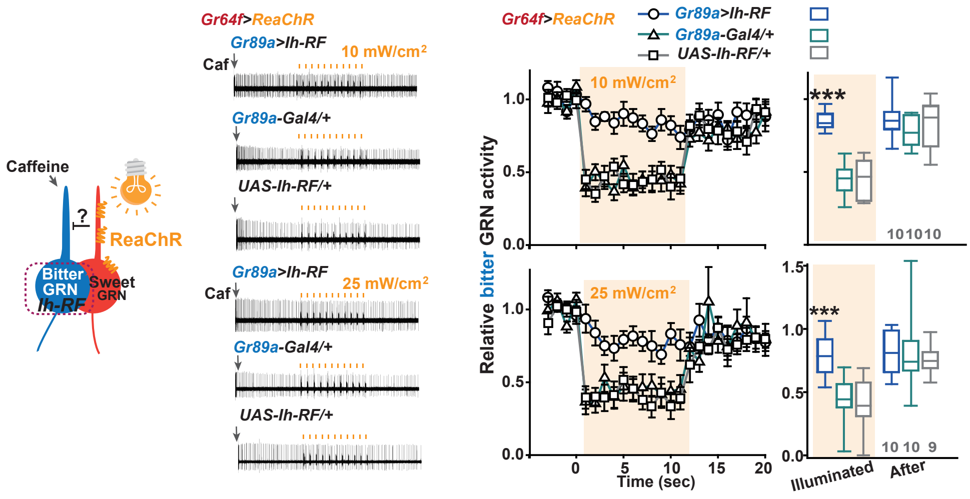


**Supplemental Fig. S6. *Ih* misexpressed in bGRNs confers strong resistance to sGRN excitationdependent inhibition.** See also Fig. 5J. At indicated irradiances, bGRNs were little inhibited by optogenetic activation of sGRNs compared to genetic controls. ***: Tukey's, $p<0.001$ between genotypes
